# Benchmarking Artificial Intelligence Models for Predicting Nuclear Receptor Activity from Tox21 Assays

**DOI:** 10.64898/2026.03.20.713297

**Authors:** Nikhil Chivukula, Jaisubasri Karthikeyan, Harshini Thangavel, Shreyes Rajan Madgaonkar, Areejit Samal

**Author notes:** N.C., J.K., H.T. and S.R.M. have contributed equally to this work and should be considered as Joint-First authors. Corresponding author: Areejit Samal, Computational Biology Group, The Institute of Mathematical Sciences (IMSc), CIT Campus, Taramani, Chennai 600113 India.

## Abstract

Tox21 assays compile extensive chemical bioactivity data across diverse biological targets, making them widely utilized resources for *in silico* model development. Nuclear receptor-specific assays within this dataset are particularly valuable for screening potential endocrine disrupting chemicals. This study presents a comprehensive benchmarking of diverse machine learning (ML), deep learning (DL), and transformer-based architectures with varied chemical feature representations across nuclear receptor assays. First, 43 datasets associated with 18 nuclear receptors within Tox21 assays were systematically curated from ToxCast invitrodb v4.3. Upon testing across these datasets, model performance was found to be dependent on the degree of class imbalance. Tree-based ML models such as random forest (RF) and extreme gradient boosting (XGBoost) trained on descriptors, or combination of descriptors and fingerprints, consistently outperformed in datasets with higher proportions of active chemicals (>10%), while DL models showed greater robustness for those with moderate proportions (5-10%). Further analysis revealed that approximately 40% of misclassified active chemicals occupied structurally isolated regions of the chemical space, suggesting absence of close structural analogues in the training set potentially contributed to their misclassification. External validation using *in vitro* and *in vivo* androgen and estrogen receptor bioactivity data showed generally good concordance. Finally, a systematic literature review revealed that the models in this study span wider range of architectures, feature representations, and assay endpoints, and are broadly comparable to or better than existing work. Overall, insights from this study can inform the development of more reliable *in silico* tools supporting new approach methodologies for nuclear receptor bioactivity predictions.

## 1. Introduction

Nuclear receptors are a class of ligand-activated transcription factors that interact with hormones, steroids, and vitamins, playing a crucial role in regulating development, reproduction, and metabolism.^1,2^ Due to their central role in such physiological processes, they are primary targets of xenobiotic endocrine disrupting chemicals (EDCs),^3^ and characterizing the interactions between environmental chemicals and nuclear receptors is essential for understanding EDC-mediated toxicities in humans. Given the large number of chemicals requiring toxicological evaluation and the limitations of traditional animal-based testing in terms of cost, time, and ethical considerations, there is a growing need for reliable *in vitro* and *in silico* approaches to screen and prioritize chemicals with potential endocrine disrupting activity.^4^ In this context, *in vitro* high-throughput screening (HTS) initiatives provide a valuable resource for the development of such predictive models.

The Toxicology in the 21^st^ Century (Tox21) program (https://tox21.gov/), a federal collaborative initiative between the United States Environmental Protection Agency (USEPA), National Institute of Environmental Health Sciences (NIEHS), National Center for Advancing Translational Sciences (NCATS), and Food and Drug Administration (FDA), has generated large-scale bioactivity data through HTS of nearly 10,000 compounds across multiple assay endpoints spanning diverse biological targets, including nuclear receptors.^5^ Since its inception, the Tox21 program has evolved across three phases, with each phase expanding both the number of chemicals screened and the breadth of assays.^5^ The program was subsequently integrated into the broader ToxCast program,^6^ where data is processed and stored in the dedicated ToxCast invitrodb^7^ database. The latest version, invitrodb v4.3 is publicly accessible,^8^ and represents one of the most comprehensive and up-to-date repositories of *in vitro* bioactivity data currently available, making it a valuable resource for the development and validation of *in silico* prediction models.

Previously, several machine learning (ML) and deep learning (DL) approaches have been applied to Tox21 data for toxicity prediction, with datasets from the Tox21 Data Challenge (https://tripod.nih.gov/tox21/challenge/) being predominantly used across published studies.^9^ However, the datasets curated for this challenge cover a limited number of nuclear receptors and do not distinguish between endpoint-specific activities such as agonism and antagonism (https://tripod.nih.gov/tox21/challenge/data.jsp), limiting their utility for mechanistically informed model development. Moreover, majority of the existing studies focus on a narrow set of model architectures and chemical representations,^10–12^ making it difficult to draw generalized conclusions about the relative strengths and limitations of different approaches across varying dataset characteristics. Recently, artificial intelligence (AI) models, especially transformer-based chemical language models have shown promise for molecular property prediction tasks,^13^ yet their performance relative to conventional ML and DL approaches under varying dataset characteristics is largely underexplored. Therefore, there is a need to systematically benchmark different AI modeling approaches on the ToxCast invitrodb nuclear receptor-specific datasets to obtain a more holistic understanding of model performance for bioactivity prediction tasks.

To address these gaps, this study extensively benchmarks ML-, DL-, and transformer-based models in combination with various chemical feature representations across a broad range of Tox21 nuclear receptor assays. In particular, the influence of class imbalance and feature representation on model performance was systematically evaluated, and the structural placement of misclassified active chemicals within the chemical similarity network (CSN) was analyzed to identify factors beyond conventional performance metrics. Additionally, external validation was further performed using *in vitro* and *in vivo* AR, ERa, and ERb bioactivity data to assess the generalizability of the developed models. Finally, a systematic literature review was conducted to contextualize the findings within existing literature, comparing modeling approaches, feature representations, and methodological choices reported therein. Overall, the findings of this study offer insights into the factors influencing model performance on Tox21 nuclear receptor bioactivity datasets, potentially contributing to the development of more reliable *in silico* approaches within new approach methodologies (NAMs).

## 2. Methods

### 2.1. Identification of nuclear hormone receptor-relevant assays within ToxCast

In this study, a systematic comparative analysis was conducted across a diverse range of artificial intelligence (AI) models, like machine learning (ML), deep learning (DL) and transformer-based models including large language models (LLMs), and feature spaces, including both structural fingerprints and molecular descriptors. This approach allowed us to evaluate the predictive utility of specific model-feature combinations and identify the most robust architectures for nuclear receptor activity classifications.

First, a list of 49 nuclear hormone receptors, along with their corresponding NCBI gene identifiers, was obtained from the HUGO Gene Nomenclature Committee (HGNC) portal (https://genenames.org/data/genegroup/#!/group/71). In addition, the Aryl Hydrocarbon Receptor (AHR) was included due to its functional similarity to nuclear receptors and its established role in reproduction, development and xenobiotic metabolism^14–17^. Next, to identify relevant Tox21 assays, the ToxCast invitrodb v4.3^6,8^ was utilized. The dataset was first filtered to obtain Tox21 assays (assays with asid 7). Next, the human-specific NCBI Gene identifiers of the 50 target receptors were cross-referenced against the target gene information for each assay to identify nuclear receptor-specific Tox21 assays. It was observed that only 18 of the 50 receptors were represented in the Tox21 assay library. Furthermore, for receptors tested in multiple assays, only the assays annotated as primary readouts relevant to agonist and antagonist conditions were retained to ensure high-confidence, mechanism-relevant chemical interactions.^18^ Through this systematic process, 30 Tox21 assays associated with 18 unique nuclear receptors were identified (Table S1).

### 2.2. Curation and standardization of chemical data within ToxCast invitrodb v4.3

#### 2.2.1. Compilation of chemical data from ToxCast invitrodb v4.3

ToxCast invitrodb v4.3 comprised 9800 chemicals tested across various assays. First, the two-dimensional (2D) structure files were obtained in v2000 SDF format from the CompTox Chemicals Dashboard^19^ (https://comptox.epa.gov/dashboard/) using their corresponding DSSTox identifiers (DTXSID). Subsequently, MayaChemTools^20^ was utilized to remove invalid structures, strip salts, and eliminate duplicated chemicals, resulting in identification of 8709 chemicals with 2D information. Thereafter, three-dimensional (3D) structures were generated from the corresponding 2D structures utilizing RDKit.^21^ The molecules were embedded in 3D space using the ETKDG method (https://www.rdkit.org/), followed by energy minimization via the MMFF94 force field. This resulted in the generation of 8554 3D chemical structures. Both 2D and 3D structures were then utilised to generate the various chemical features employed in this study.

#### 2.2.2. Calculation of chemical fingerprints and descriptors

Chemical fingerprints are machine-interpretable representations of chemical structures.^22^ In this study, RDKit (v2025.09.4) was employed to generate four types of chemical fingerprints, namely MACCS, Morgan (specifically, ECFP4 and FCFP4), and Layered fingerprints. MACCS keys were encoded as 167-bit binary vectors. Morgan fingerprints, both with features (FCFP4) and without features (ECFP4), were generated with a radius of 2 and encoded as 1024-bit binary vectors. Layered fingerprints were encoded as 2048-bit binary vectors. Further, Open Babel (v3.1.0)^23^ was utilized to obtain canonical SMILES from the 2D structures. Subsequently, 2D molecular descriptors were computed from the 2D structures, while 3D descriptors were computed from corresponding 3D structures employing both PaDEL^24^ and RDKit. A total of 1875 molecular descriptors and 231 descriptors were computed for each chemical using PaDEL and RDKit, respectively (Table S2). These descriptors included topological, geometric, electrostatic, and several other types of chemical features (Table S2). Altogether, 8430 compounds were identified for which all the features were successfully generated (Table S3). These features were then utilized in different combinations across different AI models in this study.

#### 2.2.3. Classification of receptor-specific chemical bioactivity

To define the binary activity of chemicals across the selected assays, the following consensus criteria were applied. A chemical was initially classified as ‘active’ if the associated hit-call value (hitc) was ≥ 0.9, while the rest were categorized as ‘inactive’.^7^ In cases where a chemical was associated with multiple annotations for a single assay, the chemical was considered as: (i) ‘inactive’, if there were no ‘active’ annotations across all replicates; (ii) ‘active’, if the frequency of ‘active’ annotations was greater than the frequency of ‘inactive’ annotations; (iii) ‘inconclusive’, if ‘inactive’ annotations were more frequent than, or equal to, ‘active’ annotations. The chemicals annotated as ‘inconclusive’ were excluded from further analysis to ensure high confidence chemical classification. Finally, for receptors where both agonist and antagonist data were available, a combined activity profile was created, wherein a chemical was labeled ‘active’ if it showed activity in either the agonist or antagonist assay, and labeled ‘inactive’ otherwise. Through this extensive systematic approach, the final set of chemical bioactivity labels were compiled for each of the 18 nuclear receptors (Table S3).

### 2.3. Data splitting for model training and evaluation

To train the models and evaluate their predictive performance, the dataset was partitioned using a stratified sampling approach. For traditional ML algorithms, the data was divided into an 80%-20% train-test split. For the remaining models, which require an independent set for hyperparameter tuning, the data was split 80%-10%-10% into training, validation, and test sets. It was observed that the number of ‘inactive’ compounds was much larger than the number of ‘active’ compounds in each of the nuclear receptor assays in Tox21, suggesting a class imbalance in these assays (Table 1). Thus, the stratified split function from the Python scikit-learn library^25^ was employed to ensure that the ratio of ‘active’ to ‘inactive’ chemicals remained consistent across all subsets. To further ensure the stability of the predictions and account for potential bias in data distribution, the splitting process was repeated using three different random seeds (42, 123 and 1337). Finally, the mean and standard deviations of resultant metrics were computed to get a better understanding of the model performance.

**Table 1.** List of 18 nuclear receptors, and their associated information from Tox21 assays. Note, THRA and THRB are mapped to the same assay, and therefore, have been clubbed together.

| Serial Number | Receptor | Assay | Total chemicals | Active | Inactive | % Active |
| --- | --- | --- | --- | --- | --- | --- |
| 1 | AHR | TOX21_AhR_LUC_Agonist | 7207 | 558 | 6649 | 7.74 |
| 2 | AR | TOX21_AR_BLA_Agonist_ratio | 7232 | 338 | 6894 | 4.67 |
|  |  | TOX21_AR_BLA_Antagonist_ratio | 7157 | 1074 | 6083 | 15.01 |
|  |  | Combined | 7289 | 1259 | 6030 | 17.27 |
| 3 | CAR | TOX21_CAR_LUC_Agonist | 6864 | 693 | 6171 | 10.1 |
|  |  | TOX21_CAR_LUC_Antagonist | 6873 | 526 | 6347 | 7.65 |
|  |  | Combined | 6944 | 1205 | 5739 | 17.35 |
| 4 | ERa | TOX21_ERa_BLA_Agonist_ratio | 7222 | 308 | 6914 | 4.26 |
|  |  | TOX21_ERa_BLA_Antagonist_ratio | 7193 | 714 | 6479 | 9.93 |
|  |  | Combined | 7288 | 923 | 6365 | 12.66 |
| 5 | ERb | TOX21_ERb_BLA_Agonist_ratio | 6925 | 106 | 6819 | 1.53 |
|  |  | TOX21_ERb_BLA_Antagonist_ratio | 6846 | 1169 | 5677 | 17.08 |
|  |  | Combined | 6947 | 1242 | 5705 | 17.88 |
| 6 | ESRRA | TOX21_ERR_LUC_Agonist | 6919 | 207 | 6712 | 2.99 |
|  |  | TOX21_ERR_LUC_Antagonist | 6834 | 1317 | 5517 | 19.27 |
|  |  | Combined | 6947 | 1524 | 5423 | 21.94 |
| 7 | FXR | TOX21_FXR_BLA_Agonist_ratio | 6599 | 98 | 6501 | 1.49 |
|  |  | TOX21_FXR_BLA_Antagonist_ratio | 6873 | 526 | 6347 | 7.65 |
|  |  | Combined | 6944 | 568 | 6376 | 8.18 |
| 8 | GR | TOX21_GR_BLA_Agonist_ratio | 7254 | 362 | 6892 | 4.99 |
|  |  | TOX21_GR_BLA_Antagonist_ratio | 7224 | 444 | 6780 | 6.15 |
|  |  | Combined | 7288 | 758 | 6530 | 10.4 |
| 9 | PPARd | TOX21_PPARd_BLA_Agonist_ratio | 6597 | 140 | 6457 | 2.12 |
|  |  | TOX21_PPARd_BLA_Antagonist_ratio | 6551 | 352 | 6199 | 5.37 |
|  |  | Combined | 6620 | 465 | 6155 | 7.02 |
| 10 | PPARg | TOX21_PPARg_BLA_Agonist_ratio | 7256 | 160 | 7096 | 2.21 |
|  |  | TOX21_PPARg_BLA_Antagonist_ratio | 6551 | 471 | 6080 | 7.19 |
|  |  | Combined | 7578 | 602 | 6976 | 7.94 |
| 11 | PR | TOX21_PR_BLA_Agonist_ratio | 6923 | 169 | 6754 | 2.44 |
|  |  | TOX21_PR_BLA_Antagonist_ratio | 6811 | 1676 | 5135 | 24.61 |
|  |  | Combined | 6946 | 1756 | 5190 | 25.28 |
| 12 | PXR | TOX21_PXR_LUC_Agonist | 6770 | 1917 | 4853 | 28.32 |
| 13 | RARA | TOX21_RAR_LUC_Agonist | 6589 | 200 | 6389 | 3.04 |
|  |  | TOX21_RAR_LUC_Antagonist | 6876 | 594 | 6282 | 8.64 |
|  |  | Combined | 6947 | 787 | 6160 | 11.33 |
| 14 | RORC | TOX21_RORg_LUC_CHO_Antagonist | 6868 | 548 | 6320 | 7.98 |
| 15 | RXRA | TOX21_RXR_BLA_Agonist_ratio | 6844 | 236 | 6608 | 3.45 |
| 16 | THRA/THRB | TOX21_TR_LUC_GH3_Agonist | 7292 | 24 | 7268 | 0.33 |
|  |  | TOX21_TR_LUC_GH3_Antagonist | 7182 | 1624 | 5558 | 22.61 |
|  |  | Combined | 7296 | 1643 | 5653 | 22.52 |
| 17 | VDR | TOX21_VDR_BLA_Agonist_ratio | 6615 | 49 | 6566 | 0.74 |
|  |  | TOX21_VDR_BLA_Antagonist_ratio | 6575 | 321 | 6254 | 4.88 |
|  |  | Combined | 6622 | 356 | 6266 | 5.38 |

### 2.4. Classification models for predicting nuclear receptor activity

In this study, nuclear receptor activity classification was approached through a multi-model, multi-feature framework leveraging AI, wherein traditional ML algorithms, DL architectures, and transformer-based models were assessed in combination with fingerprint-, descriptor-, and graph-based representations to identify the most predictive pairings (Figure 1).

**Figure 1.**
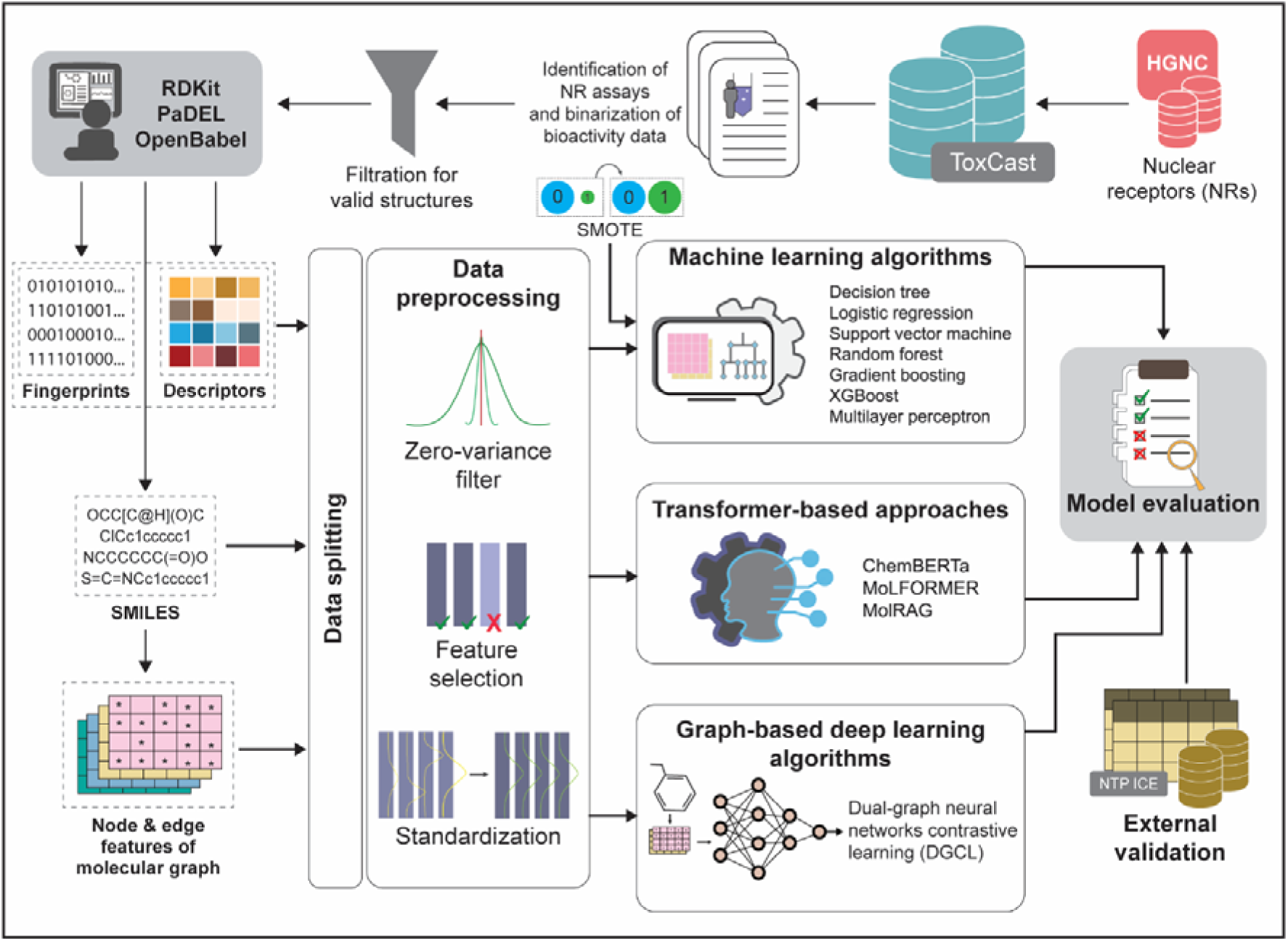
Summary of the workflow followed for the identification of nuclear receptor-specific bioactivity data from ToxCast, construction and evaluation of AI/ML-based bioactivity classification models, and evaluation of the models with external validation sets.

#### 2.4.1. Traditional machine learning algorithms

To perform the classification task, seven ML algorithms were selected from five different categories. These include a linear model (logistic regression), a tree-based model (decision tree), and three ensemble methods (random forest, gradient boosting, and extreme gradient boosting). Additionally, a kernel-based model (support vector machine) and a neural network (multi-layer perceptron) were included. For each model, three types of input representations were evaluated: (i) molecular descriptors alone; (ii) fingerprints alone; and (iii) a combination of both descriptors and fingerprints.

Logistic regression (LR) is a linear algorithm that estimates class probabilities by applying a logistic function to a linear combination of input features. The decision tree (DT) is a non-parametric algorithm that recursively partitions the feature space into hierarchical regions based on specific decision rules.^26^ To enhance stability and predictive accuracy, three ensemble methods were also employed. Random forest (RF) aggregates multiple decision trees through bootstrap sampling to reduce variance.^27^ Gradient boosting (GBT), on the other hand, builds trees sequentially to minimize a loss function by fitting to residuals.^28^ Extreme gradient boosting (XGBoost) further extends this framework using second-order derivatives in the loss optimization and regularization on the tree structure for improved performance. Additionally, the support vector machine (SVM) identifies the optimal hyperplane that maximizes the margin between classes, and uses kernel functions to handle non-linear relationships in the data.^29^ Finally, the multi-layer perceptron (MLP) is a feedforward neural network with fully connected layers and non-linear activation functions, trained via backpropagation to learn complex mappings between inputs and outputs.^30^ In this study, XGBoost was implemented using the xgboost library (https://github.com/dmlc/xgboost), while the other six models were implemented using the scikit-learn library^25^ in Python.

#### 2.4.2. Graph-based deep learning algorithm

In contrast to molecular descriptors and fingerprints, representing molecules as graphs provides an advantage as it captures the structural connectivity and spatial relationships between atoms.^31^ Graph-based representations capture atoms as nodes and chemical bonds as edges, encoding different atom and bond features within each molecule. This graph is then encoded into an embedding that can then be processed as an input by the model.^32^ In this study, Dual-graph neural networks contrastive learning (DGCL), a self-supervised graph neural network (GNN) architecture, was used for the classification tasks.^33^ Briefly, DGCL uses a two-stage learning framework where it first employs Graph Attention Network (GAT) and Graph Isomorphism Network (GIN) within a contrastive learning framework to learn features of the input molecule. These learned features are then combined with molecular fingerprints to perform the downstream prediction tasks. This approach leverages both the structural information captured by graph representations and the chemical patterns encoded in fingerprints, enabling more comprehensive molecular characterization for classification.

In this study, the DGCL architecture was modified to accommodate molecular descriptors and fingerprints independently, enabling a systematic comparison of model performance across different chemical representations. Molecular graph representations were derived from SMILES notation, and RDKit was employed to extract node and edge features that encode atomic and bond properties (Table 2). The PyTorch Geometric library^34^ was used to construct graph representations and implement the GAT and GIN architectures within DGCL.

**Table 2.** The computed node and bond features from RDKit utilized in encoding DGCL.

| Feature type | Attribute | Description | Size |
| --- | --- | --- | --- |
| Atom (Node) | Type | Atomic number of element | 44 |
|  | Degree | Number of bonds associated with the atom | 11 |
|  | Implicit Valence | Implicit valence of the atom | 11 |
|  | Formal charge | Formal charge of the atom | 1 |
|  | Number of radical electrons | Number of radical electrons of the atom | 1 |
|  | Hybridization | Hybridization associated with the atom | 6 |
|  | Aromaticity | Whether the atom is part of an aromatic ring | 1 |
|  | H atoms | Number of bonded hydrogen atoms | 11 |
| Bond (Edge) | Type | Type of bond between two atoms | 4 |
|  | Is conjugated | Whether bond belongs to a conjugated system | 1 |
|  | Is in ring | Whether bond is part of a ring structure | 1 |
|  | Stereo | Stereochemical configuration of bond | 7 |

#### 2.4.3. Transformer-based approaches

Within the AI domain, transformer-based models^35^ have emerged as powerful alternatives for molecular representation learning by treating molecular SMILES strings as sequential data. Three transformer architecture-based approaches were employed in this study, namely, ChemBERTa, MoLFORMER, and MolRAG. ChemBERTa is an encoder-based model pretrained on 100 thousand SMILES strings from ZINC database using masked language modeling, enabling it to learn robust molecular representations through self-supervised learning.^36^ MoLFORMER is a transformer model, trained on over 1.1 billion molecules from PubChem and ZINC databases, allowing it to capture structural and chemical information for diverse property prediction tasks.^37^ This study uses a variant of MoLFORMER that is trained on 10% of the data on PubChem and ZINC. MolRAG integrates retrieval-augmented generation with chain-of-thought reasoning using an LLM, such as Llama 3, as the base model.^38^ The model uses carefully designed prompts that systematically guide the language model, where each prediction is decomposed into interpretable intermediate steps that analyze structural features, functional groups, and their relationships to the target property.

SMILES representation was used as input for all the three models. The HuggingFace Transformers library (https://huggingface.co/transformers) was used to implement the ChemBERTa and MoLFORMER models. For the ChemBERTa model (https://huggingface.co/seyonec/ChemBERTa-zinc-base-v1) and the MoLFORMER model (https://huggingface.co/ibm-research/MoLFormer-XL-both-10pct), the SMILES input was tokenized into 128-bit and 256-bit vectors, respectively. A custom weighted trainer was implemented for both these models by combining class-weighted product of focal loss^39^ and cross entropy^40^. The focal loss applies a modulating factor to reduce the contribution of easily classified examples and focus on harder cases, while class weights were applied to address class imbalance by penalizing errors differently across classes. Finally, following training, the model outputs were passed through a softmax function to convert logits into predicted class probabilities for evaluation.

For MolRAG, Llama 3.1-8B instruct model obtained from Unsloth AI (https://unsloth.ai/) hosted on HuggingFace (https://huggingface.co/unsloth/Llama-3.1-8B-Instruct-GGUF) was used with llama-cpp (https://llama-cpp.com/) as the inference engine. MolRAG consisted of multiple components, namely the retrieval of molecular data for each query molecule, implementation of chain-of-thought reasoning to the language model, and unifying this combined data into a systematic prompt. First, the model was provided with general instructions regarding the prediction task. Next, dataset context was provided based on assay information from ToxCast for each of the receptors. For each query molecule in the validation set, the SMILES representations and top five descriptors identified from Boruta feature selection were retrieved for top five structurally similar reference molecules from the training set, based on Tanimoto coefficients calculated using ECFP4 fingerprints. These five neighboring molecules were further annotated with their nuclear receptor activity information to provide context to the model. The query context included the SMILES representation and descriptors for each query molecule. Finally, a molecule-specific prompt was designed to capture the dataset context, query molecule information, and information on the structurally similar reference molecules along with the task instructions. Figure 2 contains an example of the prompt utilized in MolRAG.

**Figure 2.**
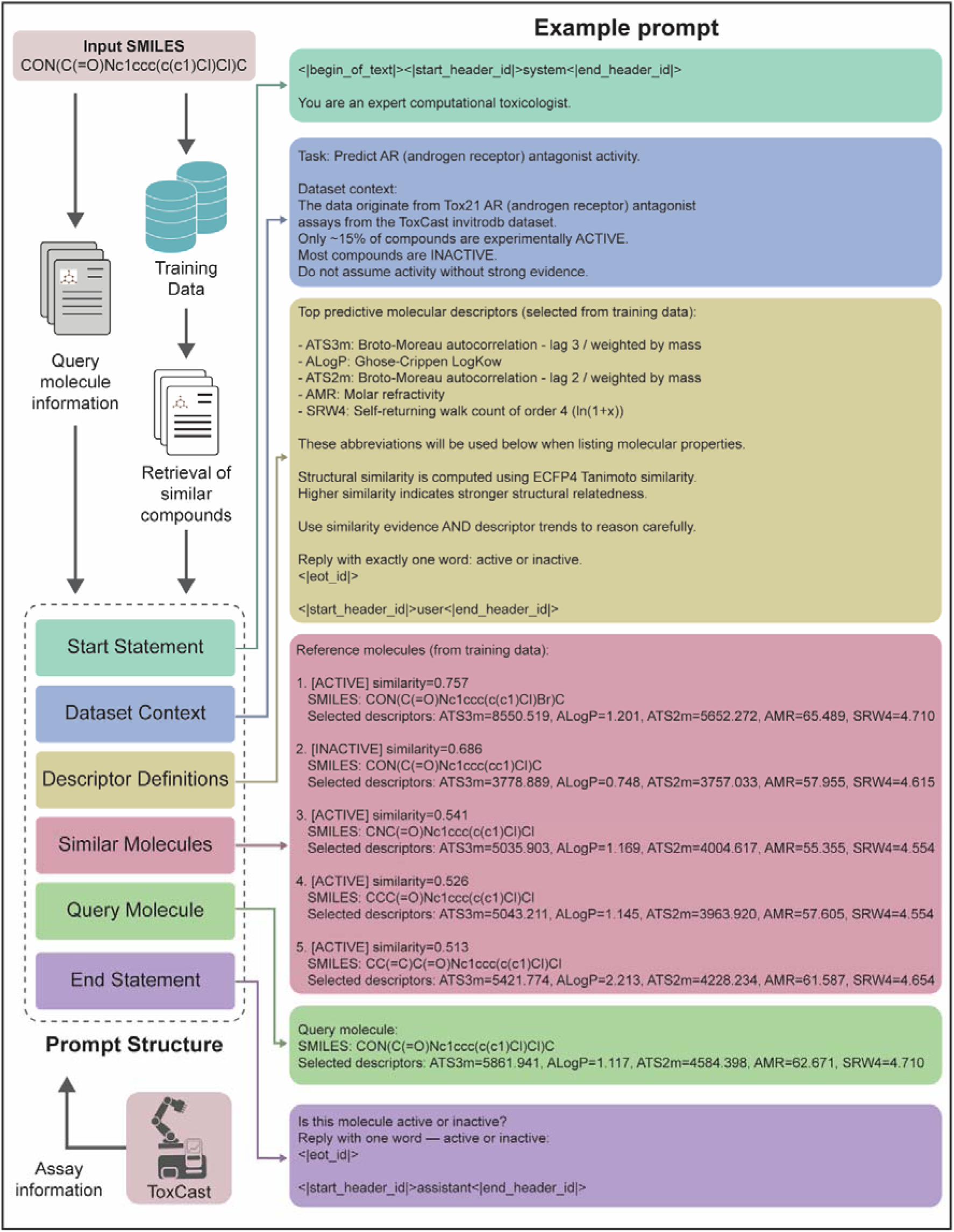
Prompt design to conduct nuclear receptor-specific bioactivity classification using MolRAG on LLama3.1-8B Instruct model.

The performance of each model was assessed using accuracy, precision, recall, F1 score, area under the Receiver Operating Characteristic curve (AUC-ROC), area under the precision-recall curve (AUC-PR), and Mathew’s correlation coefficient (MCC).

### 2.5. Data preprocessing and hyperparameter tuning

The datasets obtained from Tox21 exhibited class imbalance across all receptor assays, with varying distributions of ‘active’ and ‘inactive’ compounds. Therefore, the F1 score was selected as the primary evaluation metric as it balances precision and recall and is more suitable for imbalanced classification tasks than accuracy.^41^ Molecular descriptors and fingerprints were used as features for these algorithms to build the classification models using ML algorithms. First, for all datasets, features with zero-variance were identified and removed. Next, feature selection was performed using the Boruta algorithm implemented using boruta_py library^42^ to identify the top 100 most relevant features. Importantly, in the case of individual fingerprint data, zero-variance filtration and feature selection were not performed. A custom scaling function was designed to handle datasets containing both fingerprints and molecular descriptors, where standardization was applied only to descriptor columns while fingerprint features remained unscaled. To address class imbalance, Synthetic minority oversampling technique (SMOTE) was used to generate synthetic samples of the minority class,^43^ using three nearest neighbors to define the local neighborhood for sample generation. For the ML models, a pipeline was created consisting of scaling, SMOTE, and the classifier. The pipeline and SMOTE were implemented using the Imbalanced-learn library (https://imbalanced-learn.org) in Python. Finally, hyperparameter tuning was performed to optimize model performance across different configurations. GridSearchCV was employed with stratified 10-fold cross-validation to systematically search through predefined parameter spaces for each model. This approach ensures that each fold maintains the same class distribution as the original dataset, which is particularly important for imbalanced data.

For the DGCL model, graph representations derived from SMILES were processed directly. To address multicollinearity among descriptors, the Pearson correlation coefficient was computed for each pair of features and only one feature was retained when the correlation exceeded 0.95.^44^ For fingerprints, preprocessing involved removal of zero-variance features. For the transformer-based models, SMILES representations were used as input without preprocessing, as tokenization was handled by each model’s tokenizer. For DGCL and transformer-based models, the ROC curve was computed using the validation set, and the threshold that maximized the F1 score was identified. This optimized threshold was then applied to the test set predictions to obtain final classification results. Table S4 provides the list of hyperparameters that were tuned for each model.

### 2.6. Applicability domain analysis

The applicability domain of an *in silico* prediction model represents the chemical feature space for which the model is considered to provide reliable predictions.^45^ In this study, the Domain of Applicability (DA) index was utilized to assess the reliability of model predictions.^46–48^ The DA index is a measure based on the k-nearest neighbor (k-NN) approach using either Euclidean or Tanimoto distance, depending on the underlying feature space. The DA index for a compound is defined as the arithmetic mean of two measures, *γ* and *δ*. In this study, *γ* was computed as the mean distance of a compound to its five nearest neighbors, while *δ* was computed as the length of the mean vector to those same neighbors. For ML models relying solely on fingerprint data, the corresponding Tanimoto distances were utilized. For models using molecular descriptors, the data was first normalized, and the Euclidean distances were computed. In cases where models utilized a combination of both descriptors and fingerprints, the descriptors were normalized, and the Euclidean distances were computed for the integrated feature space. For models like DL and transformers, *γ* and *δ* values were calculated based on the Tanimoto distances between corresponding ECFP4 fingerprints. To determine the reliability of the test set predictions, the 95^th^ percentile of the DA index distribution within the training dataset was established as the threshold,^47,48^ and test compounds with a DA index within this threshold were considered to be within the model’s domain of applicability.

### 2.7. External validation

To further assess the predictive power of the models developed in this study, independent endocrine-relevant datasets from the National Toxicology Program’s (NTP) Integrated Chemical Environment (ICE) dashboard were utilized (https://ice.ntp.niehs.nih.gov/DATASETDESCRIPTION?section=Endocrine). It was observed that the dataset only provided information on agonists or antagonists of two nuclear receptors namely androgen receptor (AR) and estrogen receptor (ER). ER has two subtypes, ER alpha (ERa) and ER beta (ERb).^49^ Since the dataset did not provide subtype specific information, the ER dataset was utilized for external validation for both these receptors in this study. For each of these receptors, the *in vivo* and *in vitro* bioactivity data was obtained along with their agonist or antagonist information. Subsequently, the following systematic filtration procedure was applied to obtain the validation sets. First, the chemicals belonging to the model’s training data were excluded from the validation set. Next, it was observed that certain chemicals were labeled as both ‘active’ and ‘inactive’ in some assays, and were therefore excluded from the validation dataset. Further, the validation sets comprising 10 or fewer compounds were removed. It was observed that the validation sets for ERa and ERb agonist *in vitro* assays, and ERa and ERb antagonist assays across both *in vivo* and *in vitro* were filtered out through this systematic procedure, and were therefore not utilized for validation. Overall, this procedure resulted in identification of validation sets for AR, ERa and ERb nuclear receptors. Finally, the DA index was utilized to gauge the reliability of predictions in this external validation dataset.

### 2.8. Compilation of existing prediction models for nuclear receptors

In order to compare the performance of the models generated in this study with the existing models on nuclear receptors, a comprehensive literature review was conducted. A literature search was conducted in Scopus (https://www.scopus.com) using the following query: (TITLE-ABS-KEY (“Tox21”) OR TITLE-ABS-KEY (“ToxCast”) OR TITLE-ABS-KEY (“ToxCast bioassay”)) AND (TITLE-ABS-KEY (“machine learning”) OR TITLE-ABS-KEY (“deep learning”) OR TITLE-ABS-KEY (“artificial intelligence”)). This query was last executed on 16 March 2026 and resulted in the retrieval of 207 research articles. These were then systematically screened to identify studies focused on bioactivity classification models constructed using nuclear receptor-specific Tox21 assays (Figure S1). Thereafter, the performance of these models were compared with the models generated in this study based on computed F1 scores.

## 3. Results and Discussion

### 3.1. Identification of Tox21 assays associated with 18 nuclear receptors

Environmental chemicals, especially endocrine disrupting chemicals (EDCs), primarily target nuclear receptors, leading to a wide range of adverse effects including neurologic, developmental and reproductive disorders.^1–3^ Thus, *in silico* prediction models are constructed based on nuclear receptor bioactivity data to aid in characterization of novel EDCs.^50^ In the past, different studies have solely relied on machine learning (ML) algorithms based on chemical fingerprint data to test for nuclear receptor bioactivity classification, primarily utilizing the Tox21 dataset from ToxCast.^11,50–52^ This study aims to benchmark the performance of different AI/ML models and chemical features across nuclear receptor data from Tox21 dataset.

To construct the models, 30 Tox21 assays associated with 18 nuclear receptors were first curated from ToxCast invitrodb v4.3 (Table 1). Among these, it was observed that nuclear receptors THRA and THRB are mapped to the same assays, ‘TOX21_TR_LUC_GH3_Agonist’ and ‘TOX21_TR_LUC_GH3_Antagonist’ (Table 1), as both relate to thyroid axis disruption.^53^ Moreover, it was observed that receptors like AHR, PXR, RORC, and RXRA had either agonist or antagonist assays only as they related to corresponding toxicity mechanisms (Table 1). Specifically, AHR agonism is related to the propagation of adverse effects from environmental chemical exposure,^54^ while its inactivation is often associated with clinical treatment benefits.^55,56^ PXR is considered a master regulator of xenobiotic metabolism, and its activation is linked to the increased metabolism of hormones, which in turn alters the endocrine system,^57^ while its antagonistic effects are considered to be of therapeutic use in treatment of metabolic disorders and protecting against drug-induced toxicities.^58^ RORC antagonists can lead to toxicity through cholesterol biosynthesis, while its agonistic effects are associated with therapeutic potential in promoting inflammatory signaling.^59^ Finally, RXRA agonists promote dysfunctional adipogenesis, playing a key role in mechanisms associated with obesity,^60^ while antagonists have clinical relevance in the treatment of metabolic disorders like diabetes.^61^

In addition to these 30 assays, 13 further datasets were generated by combining both agonist and antagonist assays across receptors for which such data was available. Such aggregation will provide an overall understanding of the chemical interaction with the receptor which might be helpful in screening novel chemicals for their potential to interact with nuclear receptors.^50^

### 3.2. Exploration of chemical space of ToxCast chemicals

In this study, 8430 chemicals were systematically curated from ToxCast invitrodb v4.3, after removing salts, invalid structures and duplicated chemicals (Methods; Table S3). Thereafter, their SMILES, chemical fingerprints such as MACCS, ECFP4, FCFP4, and Layered, and descriptors such as PaDEL and RDKit, were computed (Methods). First, the chemical space of these 8430 chemicals was characterized through a chemical similarity network (CSN) constructed using Tanimoto similarity based on the ECFP4 fingerprints.^62^ When a similarity cut-off of 0.5 was applied, the CSN comprised 8430 nodes, and 15841 edges (Figure S2). The nodes were spread across 650 connected components with at least 2 chemicals each, and 2885 isolated nodes (representing 34.2% nodes) within the CSN. Among the connected nodes, the largest connected component comprised 2502 chemicals, representing 29.7% nodes within the CSN. This suggests that the ToxCast chemical space is highly diverse, consistent with its goal to maximize structural diversity of tested chemicals.^63^

Next, the chemicals and their associated bioactivity labels were systematically curated from the selected Tox21 assays (Methods). This data was then utilized to create 43 bioactivity annotated datasets associated with different combinations of receptors and activity such as agonist, antagonist and combined (Table S3), which are subsequently used to build different AI/ML models in this study. Among these datasets, it was observed that the proportion of chemicals annotated as ‘active’ was imbalanced across these assays, with the PXR agonist assay having the highest proportion of 1917 active compounds of 6770 tested chemicals (∼28.3% active ratio) and the THRA/THRB agonist assay having the lowest proportion of 24 active chemicals of 7292 tested chemicals (∼0.3%) (Table 1). The low number of active chemicals is due to highly selective behaviour associated with these receptors.^53^ Further, for each of the 43 datasets, the corresponding CSN was constructed to understand the structural distribution of chemicals annotated as ‘active’. It was observed that the active chemicals were roughly evenly distributed across the largest connected component, isolated nodes, and other connected components across these 43 datasets (Figure S3). Notably, a portion of active chemicals appeared as isolated nodes, indicating that these compounds share little structural similarity with others in the dataset. This structural uniqueness may reflect the chemical diversity of active compounds, but can also pose challenges for model training, as models may struggle to generalize structural patterns from compounds that have no similar counterparts in the dataset.^64^

### 3.3. Analysis of performance of AI/ML models

The aim of this study is to benchmark different AI/ML models against Tox21 nuclear receptor data to understand how data and its representation affect model performance. Consequently, seven ML models, namely, LR, DT, RF, GBT, XGBoost, SVM, and MLP, were constructed in combination with chemical features such as fingerprints and descriptors in different combinations (Methods). Importantly, for each of these models, a standardized pipeline was developed to eliminate data leakage issues that might otherwise lead to overestimation of the resultant classifications (Methods). This led to the generation of 49 models for each of the 43 datasets curated in this study. Next, a graph representation-based deep learning model, DGCL, was constructed wherein the inputs were chemical SMILES combined with either descriptors or fingerprints (Methods), leading to two models per dataset. Within this model, the chemical SMILES were being embedded into a graph representation, and combined with the features for training and prediction tasks (Methods). Finally, SMILES data was utilized to construct transformer-based models namely, ChemBERTa, MolFORMER, and MolRAG, leading to three additional models per dataset. Altogether, 49 ML-based models, 2 DL-based models, and 3 transformer-based models, were constructed, summing to 54 models per dataset. In each of these datasets, it was observed that the ‘active’ chemical class was present in a low proportion, resulting in a class imbalance. To address this, the F1 score was selected as the primary performance metric, as it has been shown to provide a more reliable assessment than accuracy for imbalanced datasets.^65^ Finally, the model achieving the highest average F1 score across three random splits was designated as the best-performing model for each of the 43 datasets.^66^

The reliability of *in silico* prediction models depends on their training datasets, limiting their ability to provide reliable predictions for novel chemicals.^45^ Thus, applicability domain (AD) analysis was conducted using the DA index method to filter out unreliable predictions and better gauge model performance (Methods). After AD analysis, it was observed that across all test datasets, around 3%-8% of chemicals were outside AD (Table S6). Moreover, the maximum average F1 increase was observed in the DGCL model based on descriptors for the FXR agonist dataset, indicating that prediction errors were predominantly present among the chemicals outside AD. Conversely, the maximum decrease in F1 score was observed in the MolRAG model on the PXR agonist dataset, indicating that some correct predictions were made for chemicals outside AD. Subsequently, the AD-corrected data was used to understand model performance across different datasets.

#### 3.3.1. Effect of feature type, data imbalance and underlying chemical space on model performance

Among the ML models tested across the 43 datasets, descriptor-based models had the highest average F1 score in 18 datasets, models using a combination of descriptors and fingerprints in 8 datasets, and models based on Layered fingerprints in only 2 datasets (Table S6). This suggests that ML models with descriptors, either alone or combined with fingerprints, consistently had higher average F1 scores in comparison with ML models with fingerprint-only features. It was observed that tree-based models namely RF and XGBoost were among the ML models having consistently highest average F1 scores. Furthermore, it was observed that DGCL with descriptors achieved the highest average F1 score in 9 out of 43 datasets, compared to 3 instances for DGCL with fingerprints (Table S6). Among the transformer-based models, only MoLFORMER had the highest average F1 scores, in 3 of the 43 datasets. Notably, for datasets where the active annotated chemicals constituted at least 10% of the dataset, the models trained with descriptor data, either alone or in addition to fingerprint data, consistently showed higher F1 scores across datasets (Figure 3). This is consistent with previous findings,^67^ though our results extend this observation across multiple model architectures and imbalanced datasets from ToxCast. In contrast, no clear trend was observed across models in datasets with less than 10% active compounds (Figures S3-S4).

**Figure 3.**
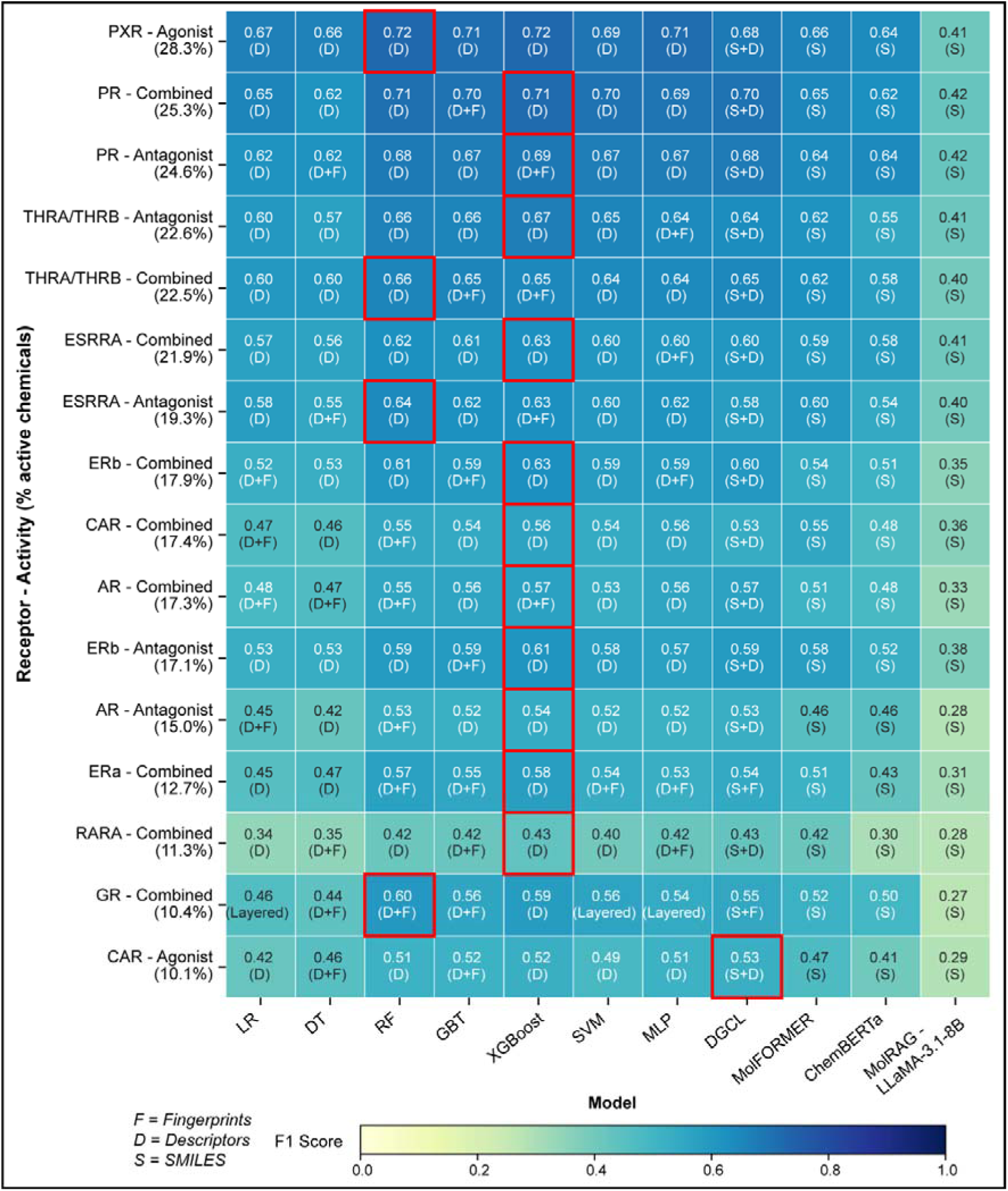
Heatmap showing the highest average F1 scores for datasets with active chemicals constituting more than 10% of total chemicals. Each row represents a dataset (receptor and activity type pair, arranged by percentage of active chemicals) and each column represents a model. Each cell displays the highest average F1 score along with the feature set used to train the model. The model with the highest average F1 score for each dataset is highlighted with a red box.

In contrast, the DGCL model based on descriptors achieved the highest average F1 scores in 6 out of 13 datasets with active compounds between 5% and 10% (Figure S4), and performed comparably to the best model in the remaining cases (Table S6). This suggests that DL-based models might have robust performance with such levels of class imbalance. However, in datasets with less than 5% active compounds, no clear trend was observed, with highly variable and dataset-dependent model-feature pairs emerging as the best performers (Figure S5), indicating that model performance in case of severe class imbalance is largely driven by the characteristics of the individual dataset.

Among transformer-based models, MoLFORMER consistently achieved higher F1 scores in comparison with ChemBERTa and MolRAG, and achieved higher average F1 scores in 3 datasets with severe class imbalance (Figure S5). MoLFORMER uses atom-wise tokenization,^37,68^ which is relatively more suited for chemical data compared to ChemBERTa, which relies on byte-pair encoding.^36^ Moreover, the underlying pre-training dataset is much larger for MoLFORMER compared to ChemBERTa, which likely contributes to its better performance.^36,37^

To understand the effect of class imbalance on model performance, the correlation between F1 scores of the model with highest average F1 score per dataset and the proportion of active compounds was evaluated using Pearson’s correlation coefficient (Figure 4). Overall, a moderate correlation was observed between these two variables (r = 0.68, *p* < 0.05). When separated by class imbalance, datasets with less than 10% active compounds showed high F1 variance and no correlation (r = 0.03, *p* > 0.05), while those with more than 10% active compounds showed a strong correlation (r = 0.85, *p* < 0.05). This suggests that when class imbalance is severe, the active ratio alone does not explain model performance, and other dataset-specific factors likely contribute to F1 variation.

**Figure 4.**
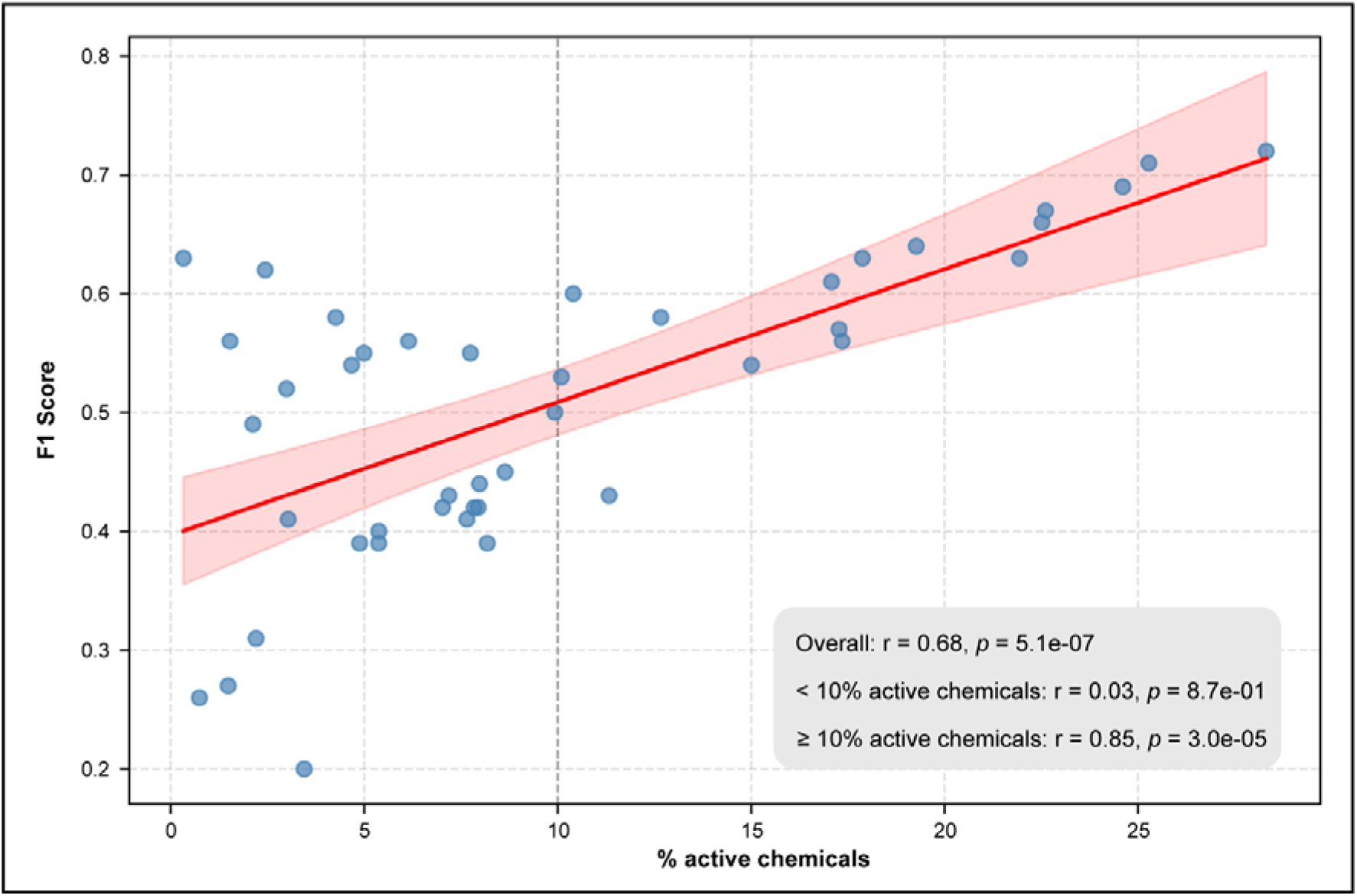
Plot depicting Pearson’s correlation between the highest average F1 score and the corresponding proportion of active chemicals (depicted as percentage of total chemicals) across 43 datasets (receptor and activity type pairs) curated in this study. The dashed vertical line at 10% divides the datasets into two regions, with Pearson’s correlation (r) calculated separately for datasets below and above this threshold, along with the overall correlation. All three values are displayed as insets within the figure.

To further explore the reason for misclassification, chemicals that were incorrectly classified as ‘inactive’ (false negatives) were mapped onto their underlying CSN (Figures S6-S8). It was observed that a notable proportion of these false negatives (∼40% on average per split) occupied isolated nodes in the CSN (Figures S6-S8). Since these structurally isolated active chemicals share no close neighbours above the similarity threshold, the model lacks the local structural context needed to learn generalizable features for the active class, likely contributing to their misclassification. This may also explain the consistently low F1 scores observed for MolRAG, as structurally isolated active chemicals would yield less informative retrieved compounds, weakening the retrieval-based reasoning context required for correct activity prediction.

#### 3.3.2. External validation of AR, ERa and ERb models

To assess the reliability and generalizability of the best models identified in this study, external validation was conducted using *in vivo* and *in vitro* endocrine disruption-relevant data obtained from the NTP ICE datasets (Methods).^69^ It was observed that this dataset provided agonist or antagonist information for only 3 of the 18 nuclear receptors considered in this study, namely AR, ERa, and ERb. Therefore, the best models based on the highest average F1 score obtained for each of these receptors, were utilized for corresponding external validation. It should be noted that the active ratio was greater than 10% across these datasets, but the number of active compounds were less as they contained between 20 and 30 chemicals (Table S7). AD analysis was conducted based on the DA index on these external datasets, and the metrics were averaged across the split-specific models.

For AR agonists, the models demonstrated reliable performance, achieving average F1 score of 0.73 for both the *in vitro* and *in vivo* datasets. Corresponding average recall values were 0.88 and 0.74, respectively, suggesting that the model was reliably predicting active chemicals within these datasets. However, in the case of AR antagonists, the *in vitro* average F1 score was 0.52 and recall was 0.71, while the *in vivo* average F1 score dropped to 0.33 and recall dropped to 0.48. This drop in performance could be attributed to the fact that the model was trained exclusively on *in vitro* data from ToxCast, and AR antagonism *in vivo* is often more complex, involving additional biological processes (e.g., metabolism) and toxicokinetics, that are not captured in *in vitro* assays.^70^

For *in vivo* ERa agonists, the best-performing model achieved an average F1 score of 0.74 and recall of 0.71. In contrast, the average F1 score dropped to 0.36 and recall to 0.24 for ERb agonists. This discrepancy can be explained by the nature of the external dataset, which was derived from rat uterotrophic assays measuring increases in uterine weight,^71^ which is primarily mediated by the ERa subtype rather than ERb.^49^ Overall, this external validation demonstrates that models generated from *in vitro* Tox21 assays can achieve decent generalizability and reliable performance on external datasets, particularly in the case of AR agonists and ERa agonists, provided the biological endpoints align.

### 3.4. Comparison of performance against published models

A systematic literature search using Scopus yielded 207 studies, of which 49 were deemed relevant based on their use of Tox21 data for specific nuclear receptor endpoints (Figure S1). Among these studies, the majority reported AUC-ROC as the primary evaluation metric, which is less informative under class imbalance conditions.^72^ Only 13 studies reported F1 score-based evaluation, and among these, considerable variation was observed in the number of assays evaluated, model architectures employed, and chemical feature representations used (Figure 5; Table S8). Of the 13 studies, 5 relied on ToxCast invitrodb, 6 relied on the Tox21 Data Challenge dataset, and 2 relied on data curated from external sources such as Collaborative Estrogen Receptor Activity Prediction Project (CERAPP)^73^ or PubChem^5^ (Figure 5; Table S8). The Tox21 Data Challenge dataset is limited to only 4 nuclear receptors and does not distinguish between agonistic and antagonistic activity, limiting its utility for mechanistic interpretation. Furthermore, this dataset is relatively dated and does not capture the full chemical set tested in later phases of the Tox21 program. Datasets derived from CERAPP and PubChem also compile data from earlier versions of Tox21 data, and are therefore unlikely to capture the most current chemical coverage. In contrast, ToxCast invitrodb is continuously updated and provides endpoint-specific activity data, making it the most comprehensive and up-to-date source for *in silico* model development.

**Figure 5:**
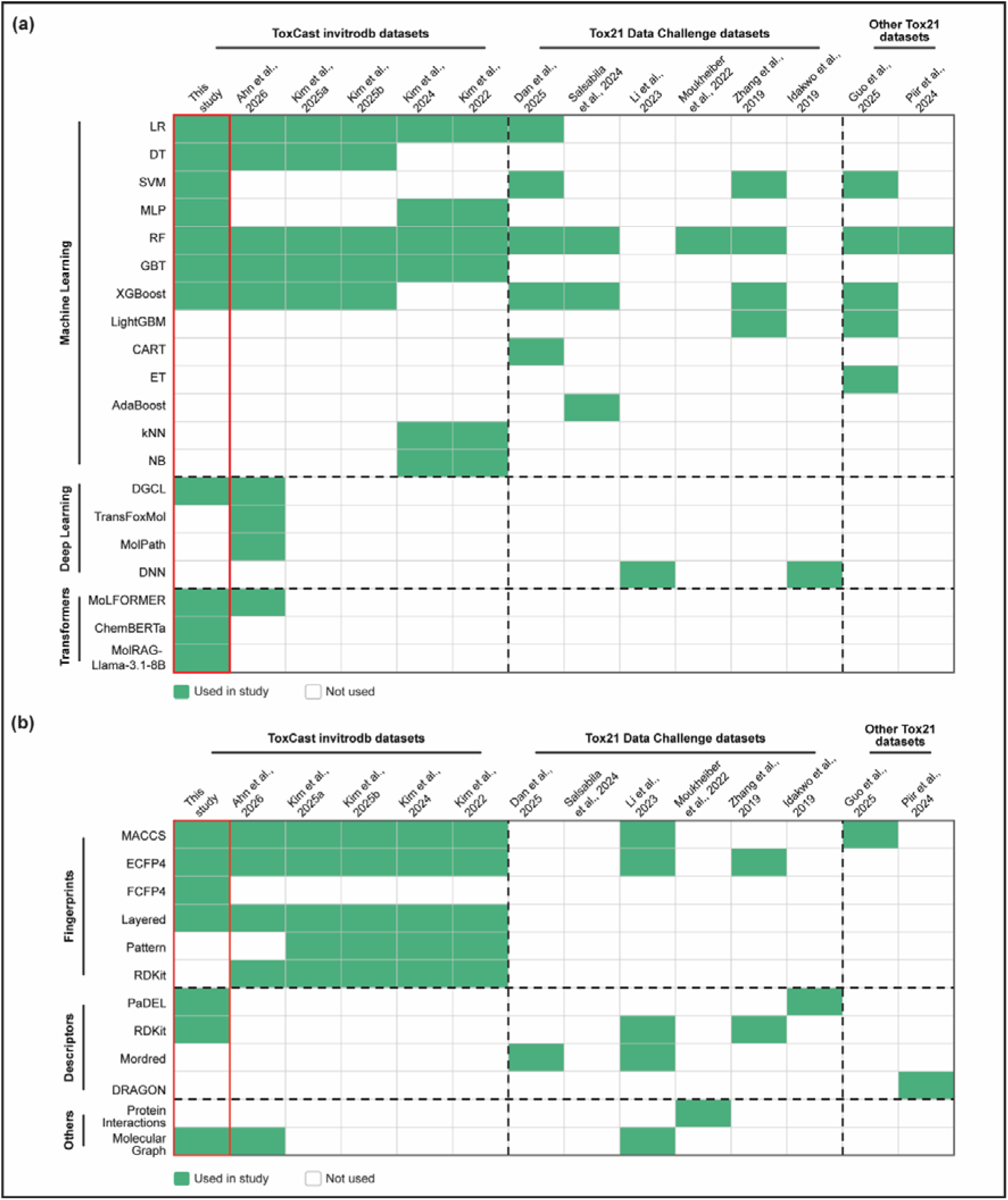
Overview of models and input features employed across curated studies wherein the studies are grouped by the dataset used for model construction. (a) Models employed across studies, categorized into machine learning (ML), deep learning (DL), and transformer-based architectures. (b) Input features used across studies, grouped into fingerprints, descriptors, and other representations. In both panels, the current study is highlighted in the red box.

Among the studies, Ahn et al., 2026 (Table S8) showed that deep learning (DL) models like DGCL consistently performed better than their machine learning (ML) counterparts on the Tox21 assays from ToxCast invitrodb, while solely relying on fingerprints. Moreover, this study covered the most assays, evaluating 28 of the 30 nuclear receptor-relevant Tox21 assays from ToxCast invitrodb. Li et al., 2023 (Table S8) showed that the inclusion of descriptor information along with fingerprint data led to improved model performance in DL models, but their study was limited to results from the Tox21 Data Challenge dataset.

In comparison, this study extensively evaluated descriptors and fingerprints, and benchmarked models beyond DL, including transformer-based models such as MolFormer, ChemBERTa, and MolRAG for 18 nuclear receptors spanning 30 Tox21 assays within ToxCast invitrodb, and additionally included combined activity profiles generated for receptors where both agonist and antagonist datasets were available. In particular, in comparison with studies performed on Tox21 assay data from ToxCast invitrodb, the current study reported higher F1 scores for the majority of the assays, and these improvements were mostly attributed to RF or XGBoost models with descriptor data alone or in combination with fingerprints. In four cases, such as PR agonist and RXRA agonist endpoints reported by Kim et al., 2024 (Table S8), and PPARd agonist and PPARg agonist endpoints reported by Kim et al., 2022 (Table S8), the previously reported F1 scores were higher than those reported in this study but still fell within the range computed in this study (Table S6).

In four other cases, such as VDR agonist and GR agonist endpoint from Kim et al., 2024 (Table S8), and AR agonist and GR agonist from Kim et al., 2022 (Table S8), the reported F1 scores were higher than the ones reported in this study. In particular, for the GR agonist endpoint, considerable differences were observed in the reported F1 scores across the two studies, with one reporting 0.85 and the other reporting 0.7. Upon closer inspection, the higher F1 score was obtained using invitrodb v2, and the lower using invitrodb v3.2 (Table S8). In both cases, F1 scores computed in this study were lower, which may be attributed to dataset differences, as invitrodb v4.3 was utilized in this study, and activity calls for the same compounds can vary across database versions.

Overall, direct comparisons across studies are constrained by differences in dataset versions, sampling strategies, and methodological choices. Nevertheless, the model performance reported in this study is broadly comparable to previously reported values, and was obtained through a more extensive and systematic evaluation across a wider range of architectures, features and assay endpoints.

## 4. Conclusion

This study systematically benchmarked different AI/ML models in combination with various chemical features across 43 nuclear receptor datasets curated from Tox21 assays. Upon curation, many datasets showed less than 10% active chemicals, with a substantial proportion of these occupying isolated nodes in the chemical similarity network (CSN), thereby posing a structural challenge to model generalizability. Among datasets with higher active ratios (>10%), ML-based models such as RF and XGBoost trained on descriptors, or a combination of descriptors and fingerprints, consistently outperformed other models. In datasets with active ratios between 5% and 10%, DL-based models showed consistently higher average F1 scores, suggesting greater robustness to class imbalance. While in datasets with less than 5% active compounds, no clear trend emerged and model performance was largely dependent on the characteristics of the individual dataset. Overall, a moderate correlation was observed between the highest average F1 score and the active ratio per dataset, whereas no such correlation was found for datasets with less than 10% active compounds, suggesting that factors beyond class imbalance influence model performance. Further analysis of false negatives revealed that approximately 40% of them occupied isolated nodes in the chemical similarity network (CSN) on average, which could contribute to limited structural context needed for proper classification. Additionally, external validation revealed good concordance for AR agonists and antagonists *in vitro*, as well as AR agonists and ERa agonists *in vivo*, while lower performance was observed for AR antagonists and ERb agonists *in vivo*, which can be attributed to complex biological mechanisms not fully captured by models trained on *in vitro* data alone. Finally, this study represents a more extensive and systematic evaluation across a wider range of architectures, feature representations, and assay endpoints, with the resulting models found to be broadly comparable to better than previously reported models.

While these findings provide useful insights, they should be interpreted in the context of different methodological constraints. The generation of descriptors relies on external tools such as PaDEL and RDKit, and any failure in descriptor computation for certain chemicals directly results in those chemicals being excluded from model training, potentially missing out on key structural information. In some cases, SVM models failed to converge on feature sets comprising descriptors and/or fingerprints in certain datasets despite extended training time, resulting in missing predictions for those splits. The performance of MolRAG may also be constrained by the use of a general-purpose language model lacking domain-specific pre-training on chemical or bioactivity data. For extremely imbalanced datasets, stratified splitting may not guarantee a meaningful distribution of active compounds across splits, suggesting the need for more appropriate splitting strategies in such cases. Although three randomized splits were generated to reduce the bias associated with the splitting strategy, this may not be sufficient to fully capture the variability in the evaluated metrics. Moreover, the high structural diversity of active chemicals in these datasets may make it difficult for models to learn generalizable structural patterns, potentially limiting their broader applicability.

Despite these limitations, this study presents a comprehensive exploration of different model architectures, including ML, DL, and transformer-based models, and chemical feature types to benchmark nuclear receptor classifications based on Tox21 data. Notably, this study observed a correlation between model performance and class imbalance across all model categories. Additionally, the analysis of chemical space topology and its relationship to model performance extends the evaluation beyond conventional performance metrics highlighting that the presence of active chemicals as isolated nodes in the CSN poses an additional challenge to model prediction, independent of class imbalance. Moreover, the inclusion of descriptor features consistently improved average F1 scores, particularly in datasets with higher active ratios. Furthermore, the integration of adverse outcome pathway (AOP) information from AOP-Wiki aids in linking model predictions to underlying biological mechanisms, providing a more interpretable framework for nuclear receptor toxicity assessment. Although, transformer-based models did not outperform descriptor-based ML and DL models in the current study, their ability to learn molecular representations directly from SMILES strings, without reliance on predefined chemical features, holds potential for capturing structural patterns beyond what conventional descriptors encode.

Overall, this study presents a systematic benchmarking of ML, DL, and transformer-based approaches across different chemical representations, evaluating their relative strengths and limitations for nuclear receptor bioactivity classification based on Tox21 assays. The insights gained from this study can aid in streamlining the development of novel *in silico* prediction models, that can contribute to the advancement of new approach methodologies (NAMs) for more reliable screening and risk assessment of potential environmental EDCs.

## Data availability

The data associated with this study is contained in the article, or in the Supplementary Information files, or in the associated GitHub repository: https://github.com/asamallab/Nuclear_Receptors-Tox21-AI.

## Acknowledgement

The authors acknowledge the High-performance computing (HPC) facility, Kamet, at The Institute of Mathematical Sciences (IMSc), Chennai, for extending the computational resources for this study. Areejit Samal would like to acknowledge funding from the Department of Atomic Energy (DAE), Government of India via Apex project to The Institute of Mathematical Sciences (IMSc), Chennai. The funders have no role in study design, data collection, data analysis, manuscript preparation or decision to publish.

## CRediT author contribution statement

**Nikhil Chivukula:** Conceptualization, Data Curation, Formal Analysis, Methodology, Software, Visualization, Writing; **Jaisubasri Karthikeyan:** Conceptualization, Data Curation, Formal Analysis, Methodology, Software, Visualization, Writing; **Harshini Thangavel:** Conceptualization, Data Curation, Formal Analysis, Methodology, Software, Visualization, Writing; **Shreyes Rajan Madgaonkar**: Conceptualization, Data Curation, Formal Analysis, Methodology, Software, Visualization, Writing; **Areejit Samal:** Conceptualization, Supervision, Formal Analysis, Methodology, Writing.

## Declaration of competing interest

The authors declare that they have no known competing financial interests or personal relationships that could have appeared to influence the work reported in this paper.

## Supplementary Figures

**Figure S1.** PRISMA diagram for curating published literature reporting nuclear receptor-specific models constructed using Tox21 assays.

**Figure S2.** Chemical similarity network (CSN) of 8430 ToxCast chemicals, which is constructed based on Tanimoto similarity between ECFP4 fingerprints of chemicals.

**Figure S3.** Stacked bar plot depicting the location of the chemicals annotated as ‘active’ on the chemical similarity network (CSN) corresponding to the 43 datasets (receptor and activity type pair) curated in this study. The 43 datasets are arranged based on their percentage of active chemicals.

**Figure S4.** Heatmap showing the highest average F1 scores for datasets with active chemicals constituting between 5% and 10% of total chemicals. Each row represents a dataset (receptor and activity type pair, arranged by percentage of active chemicals) and each column represents a model. Each cell displays the highest average F1 score along with the feature set used to train the model. The model with the highest average F1 score for each dataset is highlighted with a red box.

**Figure S5.** Heatmap showing the highest average F1 scores for datasets with active chemicals constituting below 5% of total chemicals. Each row represents a dataset (receptor and activity type pair, arranged by percentage of active chemicals) and each column represents a model. Each cell displays the highest average F1 score along with the feature set used to train the model. The model with the highest average F1 score for each dataset is highlighted with a red box.

**Figure S6.** Stacked bar plot depicting the location of the chemicals incorrectly annotated as ‘inactive’ (false negatives) on the chemical similarity network (CSN) corresponding to the 43 datasets (receptor and activity type pair) based on models trained on split 1 of the data (data stratified based on seed value 42). The 43 datasets are arranged based on the percentage of active chemicals., and the total number of incorrect predictions are mentioned within brackets.

**Figure S7.** Stacked bar plot depicting the location of the chemicals incorrectly annotated as ‘inactive’ (false negatives) on the chemical similarity network (CSN) corresponding to the 43 datasets (receptor and activity type pair) based on models trained on split 2 of the data (data stratified based on seed value 123). The 43 datasets are arranged based on the percentage of active chemicals., and the total number of incorrect predictions are mentioned within brackets.

**Figure S8.** Stacked bar plot depicting the location of the chemicals incorrectly annotated as ‘inactive’ (false negatives) on the chemical similarity network (CSN) corresponding to the 43 datasets (receptor and activity type pair) based on models trained on split 3 of the data (data stratified based on seed value 1337). The 43 datasets are arranged based on the percentage of active chemicals., and the total number of incorrect predictions are mentioned within brackets.

## Supplementary Tables

**Table S1:** This table contains the information on the 30 Tox21 assays that are associated with 18 nuclear receptors. For each assay, the table provides the corresponding endpoint identifier, name, description, function type, data type, key positive control, signal direction, intended target type and sub-type, intended target family and sub-family information curated from ToxCast invitrodb v4.3. In addition, for each assay, the table provides the corresponding target gene information like entrez identifier(s) (separated by ‘|’ symbol), name(s) (separated by ‘|’), official symbol(s) (separated by ‘|’ symbol), taxonomic applicability, and associated information on molecular initiating events (MIE) from AOP-Wiki, and the associated adverse outcome pathway (AOP) (separated by ‘|’ symbol).

**Table S2:** This table contains the list of 1875 PaDEL and 231 RDKit descriptors computed in this study. For each descriptor, the table provides the descriptor name, the description, 2D or 3D class categorization of the descriptor, and the tool from which the descriptor was obtained. Note, the descriptor TPSA is computed in both the tools, and the descriptor in RDKit has been renamed to RDKit_TPSA to remove redundancy.

**Table S3:** This table contains information on the 8430 chemicals curated from ToxCast invitrodb v4.3. For each chemical, the table provides the corresponding DSSTox identifier, Chemical Abstracts Service Registry Number (CASRN), and canonical SMILES. Additionally, for each chemical, this table provides the activity information in the curated Tox21 assays from Table S1. Here ‘0’ represents ‘inactive’ and ‘1’ represents ‘active’.

**Table S4:** This table provides the hyperparameters utilized to tune the models. For each model the table provides the name, category of the model, corresponding hyperparameters, and the value(s) used.

**Table S5:** This table contains the computed metrics of all the models. For each model, the table provides the receptor, activity type, name of model evaluated, the corresponding feature set used for training the model, the threshold value, accuracy, precision, recall, F1 score, MCC, AUC-ROC, and AUC-PR. This data is mentioned across all 3 random splits utilized in this study. Finally, the average values of the metrics such as accuracy, precision, recall, F1 score, MCC, AUC-ROC, and AUC-PR are provided with the corresponding standard deviation mentioned within brackets.

**Table S6:** This table contains the computed metrics of all the models after applicability domain analysis. For each model, the table provides the receptor, activity type, name of model evaluated, the corresponding feature set used for computing applicability domain of the model, the DA index threshold value corresponding to 95th percentile in train set, total number of test chemicals, number of test chemicals within the domain of the model, accuracy, precision, recall, F1 score, MCC, AUC-ROC, and AUC-PR. This data is mentioned across all 3 random splits utilized in this study. Finally, the average values of the metrics such as accuracy, precision, recall, F1 score, MCC, AUC-ROC, and AUC-PR are provided with the corresponding standard deviation mentioned within brackets.

**Table S7:** This table provides the computed model metrics after evaluation based on external validation sets of AR, ERa and ERb. These evaluation metrics were calculated specifically for the chemicals remaining after the applicability domain analysis. For each receptor and activity type, the table provides the name of model evaluated, the corresponding feature set used for training the model, number of chemicals in the set, number of active chemicals, total number of chemicals within domain, accuracy, precision, recall, F1 score, MCC, AUC-ROC, and AUC-PR. This data is mentioned across models generated from all 3 random splits utilized in this study. Finally, the average values of the metrics such as accuracy, precision, recall, F1 score, MCC, AUC-ROC, and AUC-PR are provided with the corresponding standard deviation mentioned within brackets.

**Table S8:** This table provides the curated studies pertaining to models constructed for nuclear receptors utilizing Tox21 data. For each study, the table provides the assay information such as the number of assays tested, the receptor - activity in the assay and the assay name, information on features such as the number and types tested and their name(s) (separated by ‘, ‘), the dataset split for train, test and validation sets within the study, model architecture information such as number and architectures tested and their name(s) (separated by ‘, ‘), best performing model information such as the architecture and feature corresponding to the assay, model performance metrics such as the F1 score, accuracy, precision, recall, MCC, AUC-ROC, and AUC-PR, and the reference to the study denoted by DOIs.

## Notes

### Competing Interest Statement

The authors have declared no competing interest.

