## Supplementary material for "Benchmarking Artificial Intelligence Models for Predicting Nuclear Receptor Activity from Tox21 Assays": Figure S

### for

### Supplementary Figures

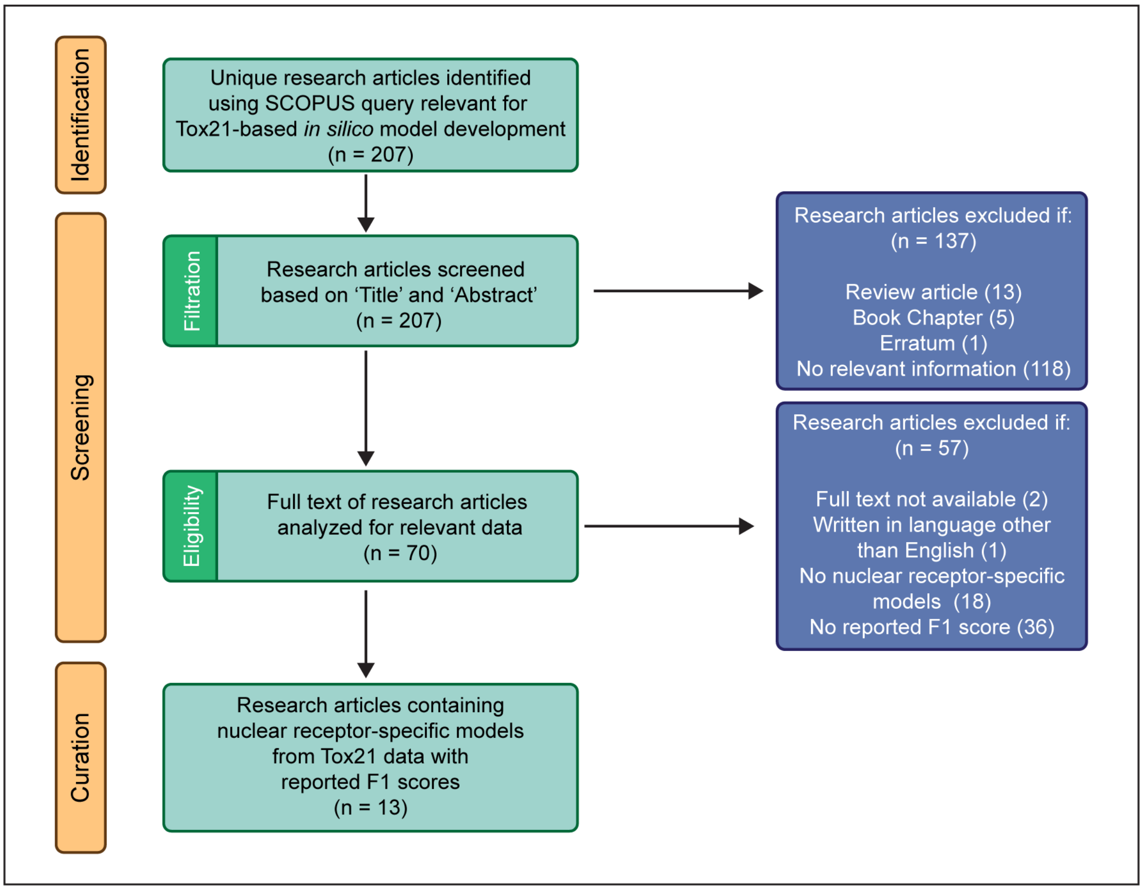


**Figure S1.** PRISMA diagram for curating published literature reporting nuclear receptor-specific models constructed using Tox21 assays.

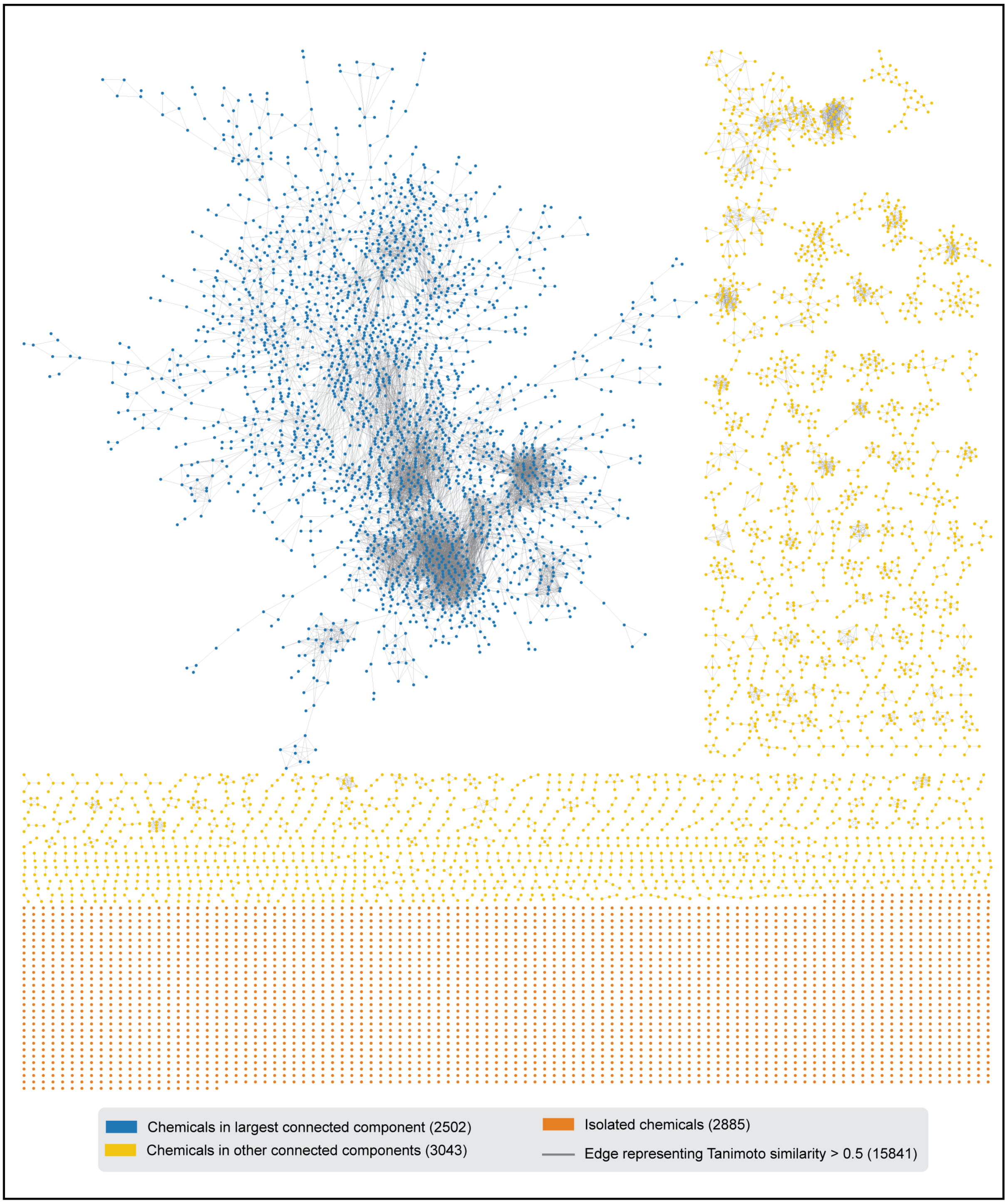


**Figure S2.** Chemical similarity network (CSN) of 8430 ToxCast chemicals, which is constructed based on Tanimoto similarity between ECFP4 fingerprints of chemicals.

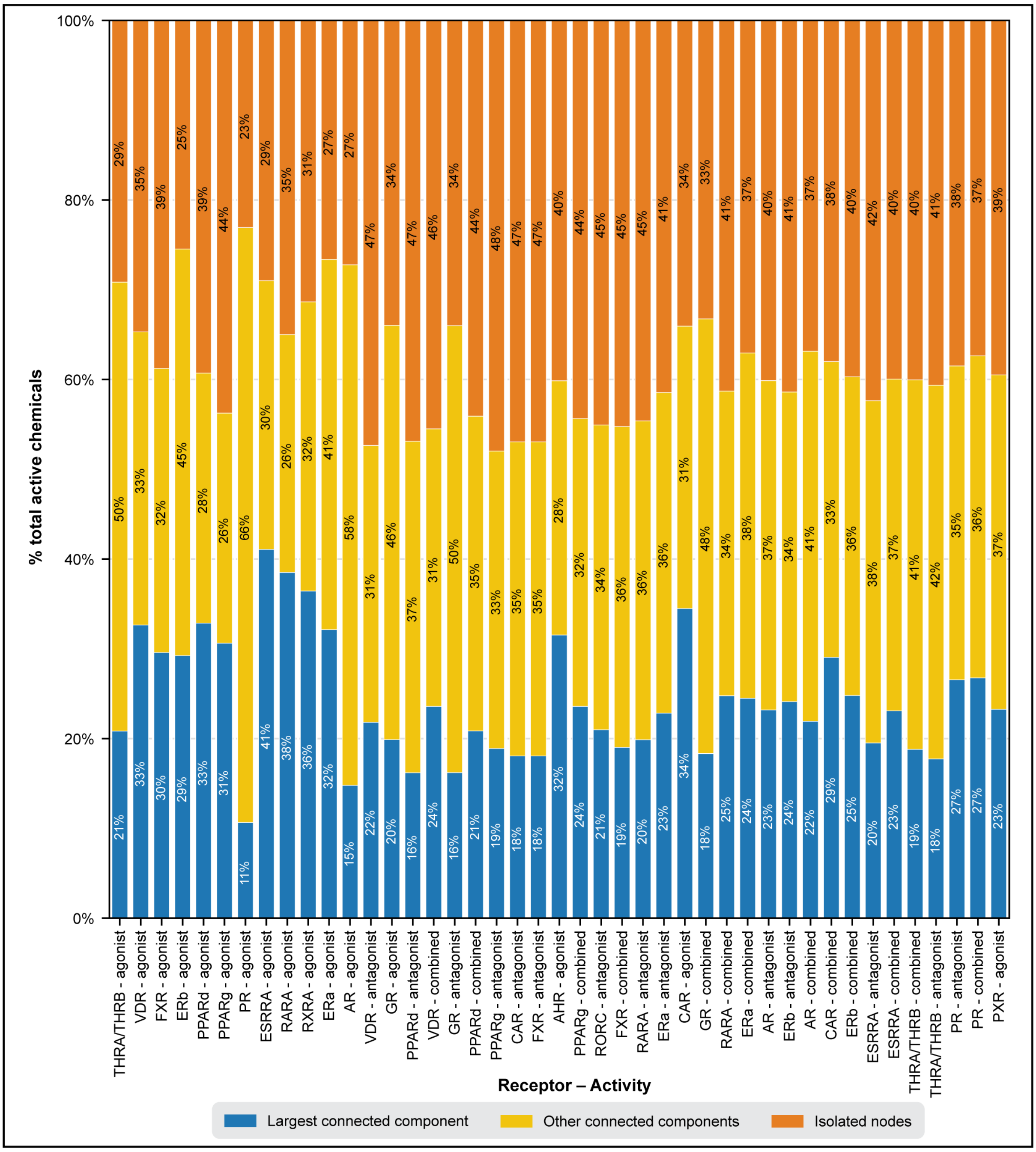


**Figure S3.** Stacked bar plot depicting the location of the chemicals annotated as 'active' on the chemical similarity network (CSN) corresponding to the 43 datasets (receptor and activity type pair) curated in this study. The 43 datasets are arranged based on their percentage of active chemicals.

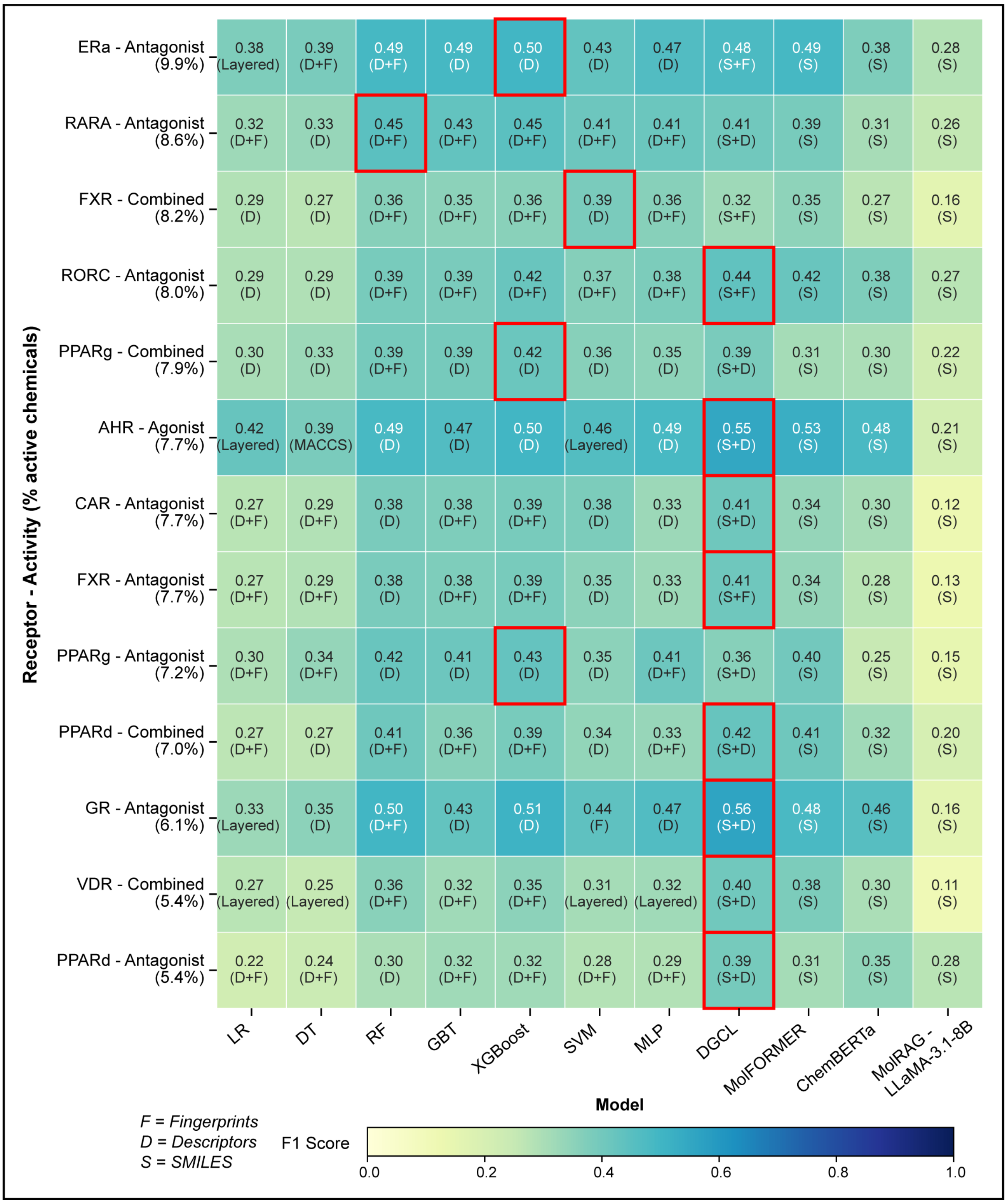


**Figure S4.** Heatmap showing the highest average F1 scores for datasets with active chemicals constituting between 5% and 10% of total chemicals. Each row represents a dataset (receptor and activity type pair, arranged by percentage of active chemicals) and each column represents a model. Each cell displays the highest average F1 score along with the feature set used to train the model. The model with the highest average F1 score for each dataset is highlighted with a red box.

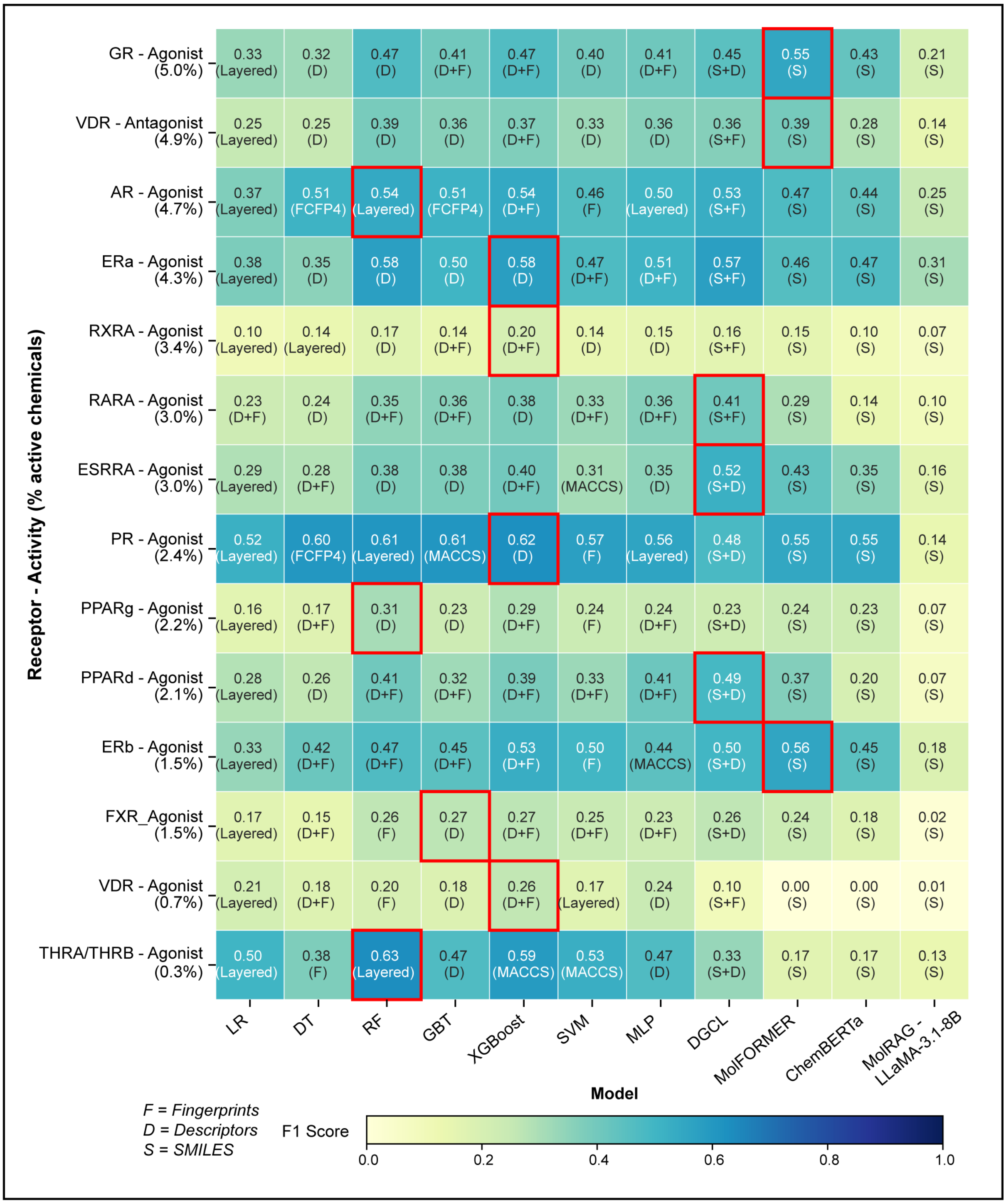


**Figure S5.** Heatmap showing the highest average F1 scores for datasets with active chemicals constituting below 5% of total chemicals. Each row represents a dataset (receptor and activity type pair, arranged by percentage of active chemicals) and each column represents a model. Each cell displays the highest average F1 score along with the feature set used to train the model. The model with the highest average F1 score for each dataset is highlighted with a red box.

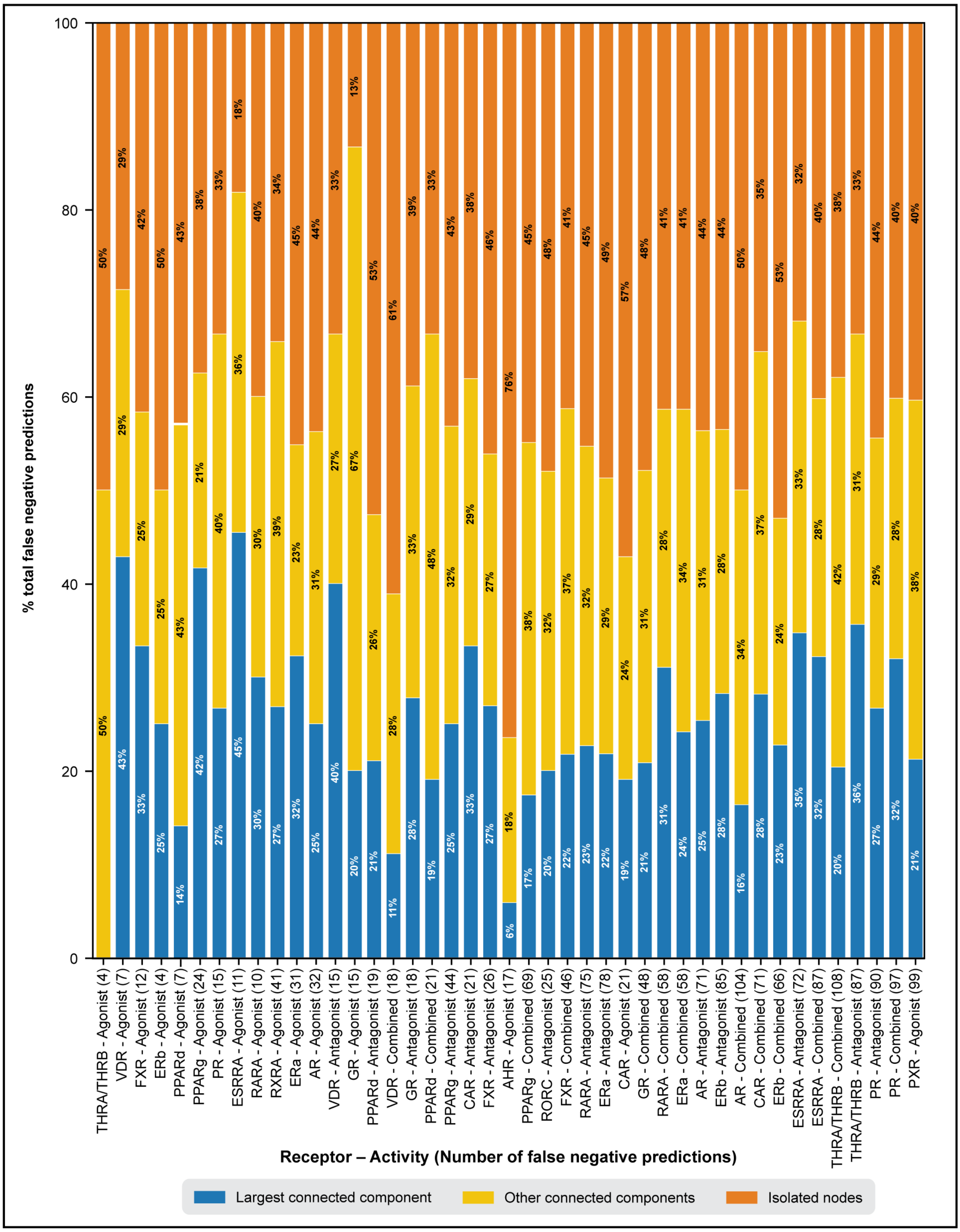


**Figure S6.** Stacked bar plot depicting the location of the chemicals incorrectly annotated as 'inactive' (false negatives) on the chemical similarity network (CSN) corresponding to the 43 datasets (receptor and activity type pair) based on models trained on split 1 of the data (data stratified based on seed value 42). The 43 datasets are arranged based on the percentage of active chemicals., and the total number of incorrect predictions are mentioned within brackets.

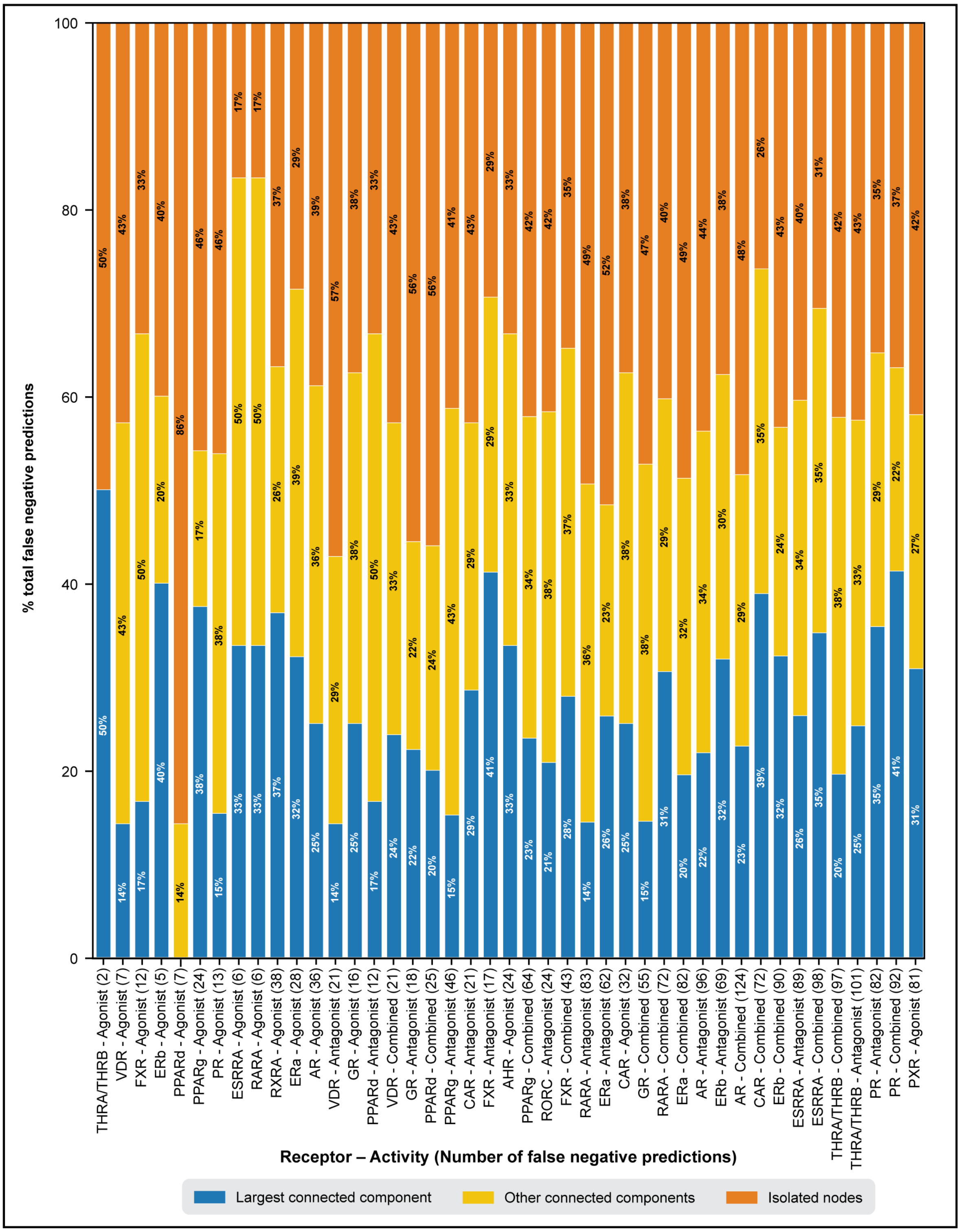


**Figure S7.** Stacked bar plot depicting the location of the chemicals incorrectly annotated as 'inactive' (false negatives) on the chemical similarity network (CSN) corresponding to the 43 datasets (receptor and activity type pair) based on models trained on split 2 of the data (data stratified based on seed value 123). The 43 datasets are arranged based on the percentage of active chemicals., and the total number of incorrect predictions are mentioned within brackets.

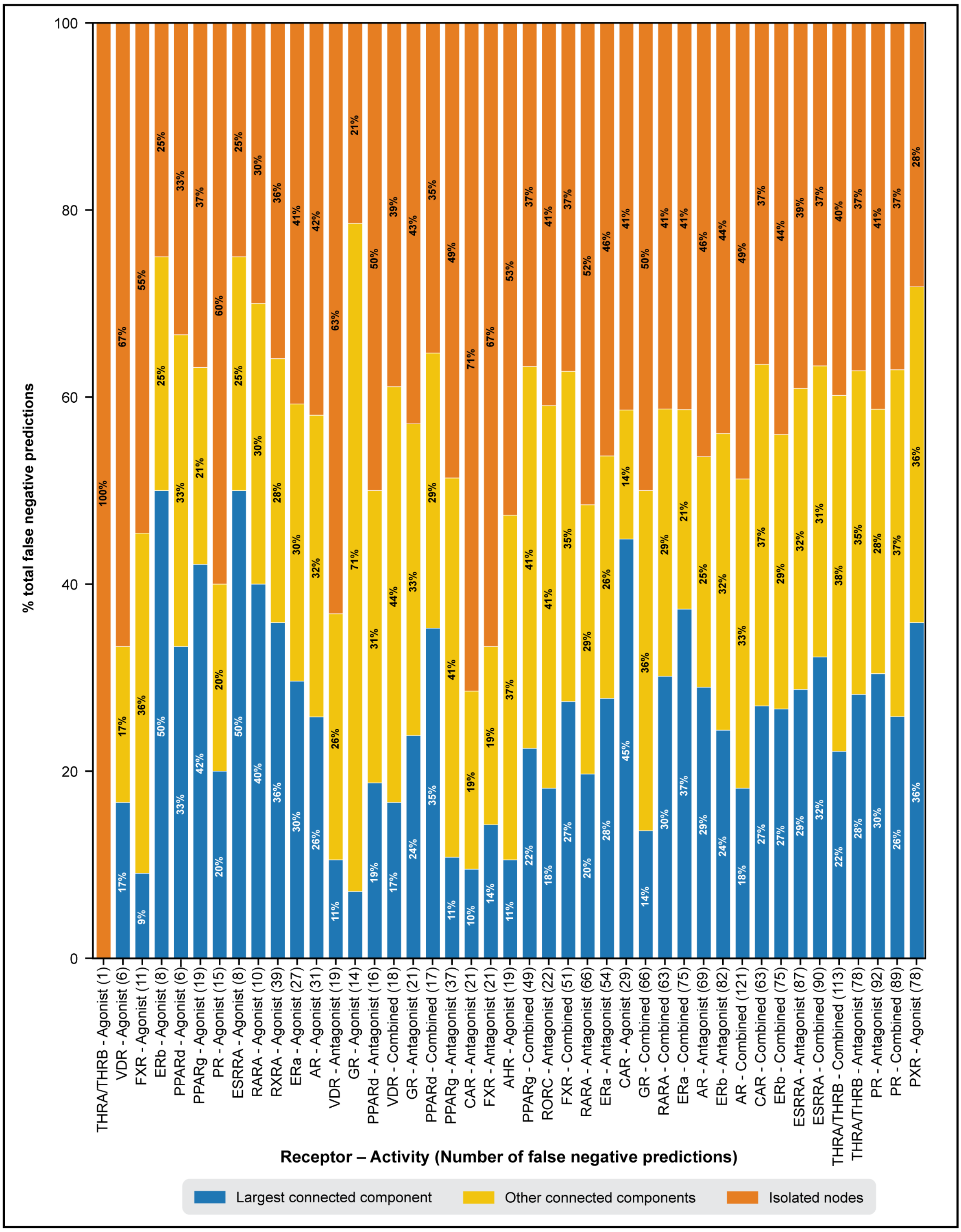


**Figure S8.** Stacked bar plot depicting the location of the chemicals incorrectly annotated as 'inactive' (false negatives) on the chemical similarity network (CSN) corresponding to the 43 datasets (receptor and activity type pair) based on models trained on split 3 of the data (data stratified based on seed value 1337). The 43 datasets are arranged based on the percentage of active chemicals., and the total number of incorrect predictions are mentioned within brackets.
